# scCausalVI disentangles single-cell perturbation responses with causality-aware generative model

**DOI:** 10.1101/2025.02.02.636136

**Authors:** Shaokun An, Jae-Won Cho, Kai Cao, Jiankang Xiong, Martin Hemberg, Lin Wan

## Abstract

Single-cell RNA sequencing provides detailed insights into cellular heterogeneity and responses to external stimuli. However, distinguishing inherent cellular variation from extrinsic effects induced by external stimuli remains a major analytical challenge. Here, we present scCausalVI, a causality-aware generative model designed to disentangle these sources of variation. scCausalVI decouples intrinsic cellular states from treatment effects through a deep structural causal network that explicitly models the causal mechanisms governing cell-state-specific responses to external perturbations while accounting for technical variations. Our model integrates structural causal modeling with cross-condition in silico prediction to infer gene expression profiles under hypothetical scenarios. Comprehensive benchmarking demonstrates that scCausalVI outperforms existing methods in disentangling causal relationships, quantifying treatment effects, generalizing to unseen cell types, and separating biological signals from technical variation in multi-source data integration. Applied to COVID-19 datasets, scCausalVI effectively identifies treatment-responsive populations and delineates molecular signatures of cellular susceptibility.

**Code availability:** Software is available at https://github.com/ShaokunAn/scCausalVI.

## 1 Introduction

Single-cell RNA sequencing (scRNA-seq) provides unprecedented insights into cellular heterogeneity and the molecular mechanisms governing cell fate and function. By profiling gene expression at the individual cell level, scRNA-seq uncovers cellular subpopulations, differentiation pathways, and responses to stimuli often masked in bulk analyses. The most common experimental paradigm is case-control studies which make it possible to understand how cells respond to perturbations such as genetic modifications [1], drug treatments [2], or environmental changes [3]. Using scRNA-seq as a readout, case-control studies facilitate the dissection of complex biological processes by revealing how individual cells respond to external stimuli [4].

However, analyzing perturbed or treated single-cell data presents significant computational challenges. Individual cells cannot be measured simultaneously across multiple experimental conditions due to cellular destruction during RNA extraction, preventing direct comparison of cellular states between conditions. Besides, observed differences between unpaired cells in control and treated conditions are the outcome of both inherent cellular heterogeneity and treatment effects. This entanglement complicates the attribution of observed transcriptional changes to specific perturbations versus pre-existing cellular states. Moreover, intrinsic cell states and phenotypic variations modulate cellular responses to treatments, further complicating the task of disentangling cell-state-specific effects from inherent cellular heterogeneity [5, 6].

To investigate the underlying treatment effects in single-cell data, several computational methods have been developed to quantify cellular responses in gene expression space. Generative models predict cellular responses to perturbations primarily at the population level, and they implicitly assume homogeneous cell responses [7, 8, 9, 10, 11]. Disentanglement methods use contrastive learning to identify salient features of treated cells through latent representations [12, 13, 14]. However, they assume independence between cellular identity and treatment effect factors, and consequently, they fail to capture the complex interplay between sources of variations [15]. Optimal transport-based approaches align cellular distributions across conditions to create pseudo-pairings for studying perturbational effects [16, 17]. While they account for distributional shifts, they do not explicitly model how intrinsic cellular heterogeneity leads to differential treatment responses. In conclusion, these methods have limited ability to capture cell-specific variations in treatment response that arise from complex cellular mechanisms including off-target effects [18] and state-dependent responses [5].

Causal inference provides a robust framework to address these limitations by modeling underlying causal mechanisms and systematically accounting for confounding factors [19, 20, 21]. By distinguishing causation from mere correlation, causal inference can identify intrinsic treatment effects and disentangle intertwined sources of variation. Structural causal models (SCMs) [22], in particular, enable the decomposition of observed variations into interpretable components, enhancing both the model’s interpretability and generalizability. Besides, SCMs facilitate interventions, allowing researchers to simulate the effects by manipulating certain variables and predicting outcomes under hypothetical scenarios [23, 24]. These methodologies have been employed to model scRNA-seq data, facilitating more precise interpretations and uncovering underlying biological mechanisms [25, 26, 16, 27, 28].

Building upon these insights, we propose scCausalVI, a causality-aware generative model designed to disentangle inherent cellular heterogeneity from differential treatment effects at the single-cell level, particularly in the context of case-control studies. By encoding the principle of SCM into the architecture of deep neural networks, the contributions of scCausalVI are twofold. First, scCausalVI effectively disentangles and explicitly models the causal relationships between inherent cellular states and treatment effects with distinct sets of latent variables [15]. The SCM framework allows for a precise characterization of how inherent cellular heterogeneity modulates treatment-specific responses. Second, scCausalVI utilizes the SCM to perform cross-condition in silico prediction, where cellular states are computationally predicted under alternative experimental conditions. This enables systematic comparison of cellular states across conditions for individual cells, predicting how the gene expression profile of an untreated cell would evolve under treatment conditions. By enabling these computational perturbation analyses at single-cell resolution, scCausalVI aids in the identification of key regulatory mechanisms and potential therapeutic targets [29].

We demonstrate scCausalVI’s effectiveness by (1) outperforming existing methods in disentangling cellular heterogeneity from treatment effects on simulated data; (2) capturing diverse immune cell responses and demonstrating robust generalization on interferon-*β* (IFN-*β*) stimulated peripheral blood mononuclear cell (PBMC) data; (3) simultaneously disentangling intrinsic cellular states, treatment effects, and batch effects across independent COVID-19 PBMC datasets, which were further validated through negative control experiments; (4) identifying treatment-responsive populations in respiratory epithelial cells through cross-condition in silico prediction; and (5) revealing transcriptional signatures that distinguish resistant from susceptible phenotypes in COVID-19 PBMC analysis.

## 2 Results

### 2.1 Overview of scCausalVI framework

scRNA-seq has emerged as a powerful tool for understanding cellular responses to perturbations, particularly in case-control studies. In these experimental designs, observed transcriptional variations arise from two sources: intrinsic cellular heterogeneity and treatment-induced effects (Fig. 1a). To deconvolute these intertwined sources of variation, we propose scCausalVI, a causality-aware deep learning framework that enables causal inference of treatment effects at single-cell resolution.

**Figure 1:**
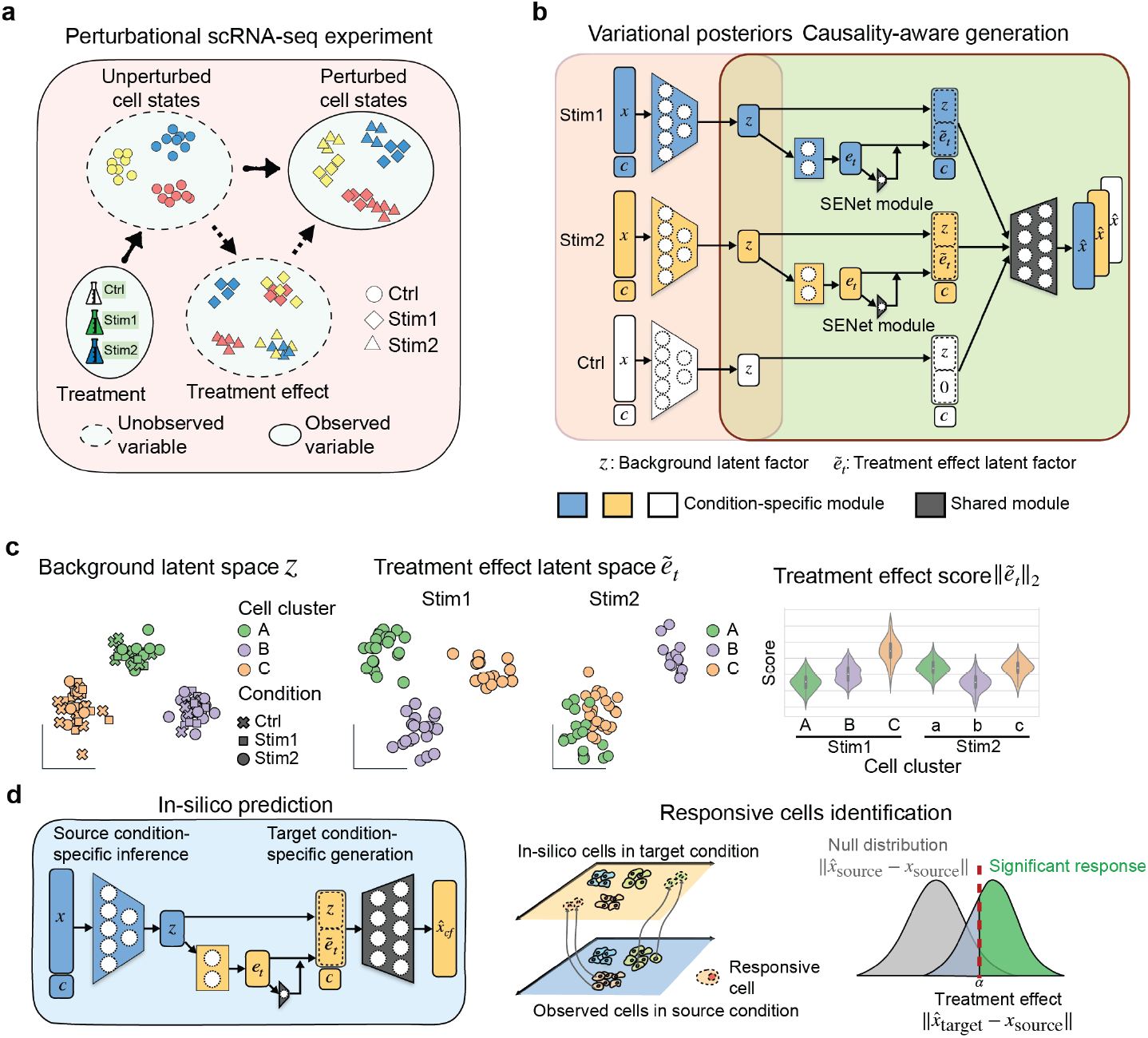
Overview of scCausalVI. **a**, Schematic of a perturbational scRNA-seq experiment, illustrating the interplay between cellular baseline states and cell-state-specific treatment effects. **b**, The causality-aware neural network of scCausalVI, consisting of variational inference through condition-specific encoders, and causality-aware generation with SCM featuring SENet attention modules for adaptive scaling, and shared decoding of gene expression profiles. **c**, Downstream analyses using latent representations to reveal inherent cellular heterogeneity pattern in background latent space *z*, and differential response in treatment effect latent space 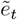. The treatment effect score, *L*2-norm of 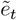, is used to quantify the treatment effect size. **d**, Cross-condition in silico prediction pipeline that predicts gene expression profiles under target conditions for cells observed in source conditions. Responsive cells can be identified by quantifying the significance of the induced difference between observed and cross-condition predicted cellular states.

scCausalVI analyzes case-control scRNA-seq data from randomized experimental designs, where it is assumed that cells are allocated to control and treatment conditions in an unbiased manner. This experimental framework enables unbiased estimation of causal effects by controlling for pre-existing cellular differences [21]. It advances beyond existing approaches by integrating SCM with variational inference to learn causal generation with unobserved factors (Fig. 1b). scCausalVI learns two distinct but interrelated sets of latent variables: background factors capturing inherent cellular states and treatment effect factors encoding treatment-induced transcriptional changes. Through a Squeeze-and-Excitation Networks (SENet) attention mechanism [30], the model adaptively scales treatment effects for individual cells, enabling cell-state-specific response modeling at single-cell resolution. This framework enables the disentangled characterization of baseline cellular heterogeneity and treatment responses while preserving their mechanistic dependencies through the SCM structure. When batch information is available, scCausalVI can additionally account for technical variations by incorporating batch indices in its inference and generation modules, enabling the elimination of technical batch effects from biological variation in both the background and treatment effect latent spaces. This comprehensive framework thus provides a principled approach for dissecting the complex interplay between cellular states, treatment responses, and batch effects in single-cell studies.

The learned latent representations from scCausalVI facilitate comprehensive biological insights through multiple analytical approaches (Fig. 1c). The background latent space reveals cellular heterogeneity patterns independent of treatment effects, while the treatment effect space uncovers subpopulations with shared response characteristics. Quantitatively, an *L*2-norm of the treatment effect latent factors serves as a measure of effect size at single-cell resolution, facilitating the identification of differential responsive patterns within treated populations.

A distinctive feature of scCausalVI is its ability to perform in silico perturbation at single-cell resolution. By intervening in condition settings within SCM, scCausalVI predicts gene expression profiles under hypothetical scenarios. We define treatment-responsive cells as those exhibiting measurable transcriptional changes in response to treatment. To identify these cells, scCausalVI generates both factual (same condition) and cross-condition (alternative target condition) predictions for each observed cell in the source condition (Fig. 1d). The difference between the cross-condition predictions and observations indicates the treatment-induced changes while controlling for intrinsic cellular heterogeneity. These cells are identified by comparing treatment-induced differences against a null distribution of generative uncertainty, which is quantified by the differences between observations and their factual predictions. Subsequently, differential gene expression analysis between responsive and non-responsive cohorts enables molecular characterization of treatment-induced biological changes.

### 2.2 scCausalVI revealed cell-state-specific treatment effects on simulated data

We compared scCausalVI with state-of-the-art methods for treatment effects estimation, including the disentangled learning models contrastiveVI [12] and scDisInFact [14], the causal-inference-based model CINEMA-OT [16], and the cell-attribute-aware models scGen [7], CPA [8], and biolord [10]. We highlight that scCausalVI requires only condition labels, operating without supervision from cell attribute information. This unsupervised strategy helps mitigate potential biases arising from cell type annotations, which can be particularly problematic when perturbations or treatments alter the expression of marker genes commonly used for cell type identification [31]. A comprehensive comparison of the features and capabilities of these methods is provided in Supplementary Table 1.

**Table 1:** Comparison of different methods and their capabilities.

|  | disentangled representation | Cell-state-specific treatment effect | cross-condition prediction | cell-type -free | generalization to unseen cells |
| --- | --- | --- | --- | --- | --- |
| scCausalVI | Yes | Yes | Yes | Yes | Yes |
| contrastiveVI | Yes | No | Yes | Yes | Yes |
| CINEMA-OT | Yes | Yes | No | Yes | No |
| scDisInFact | Yes | No | Yes | Yes | Yes |
| CPA | Yes | No | Yes | No | Yes |
| scGen | No | No | Yes | No | Yes |
| biolord | Yes | No | Yes | No | Yes |

We evaluated the different models using a simulated scRNA-seq dataset. We generated realistic synthetic data with scDesign3 [32] based on an IFN-*β*-stimulated scRNA-seq data [33], comprising four cell types across a control group and two treated groups (Fig. 2a). In each treated group, one cell type was exclusively perturbed, resulting in six distinct cellular populations under three conditions (Fig. 2b). We first assessed the prediction accuracy of scCausalVI and other generative methods by comparing the observed and predicted data (Supplementary Fig. 1). scCausalVI demonstrated exceptional performance, with predicted data aligning perfectly with the observations in UMAP space and exhibiting a near-perfect correlation (*R*^2^ *≈* 1.00) when using mean gene expression across all features. While scGen and biolord also showed high reconstruction accuracy, contrastiveVI, scDisInFact, and CPA did not perform as well.

**Figure 2:**
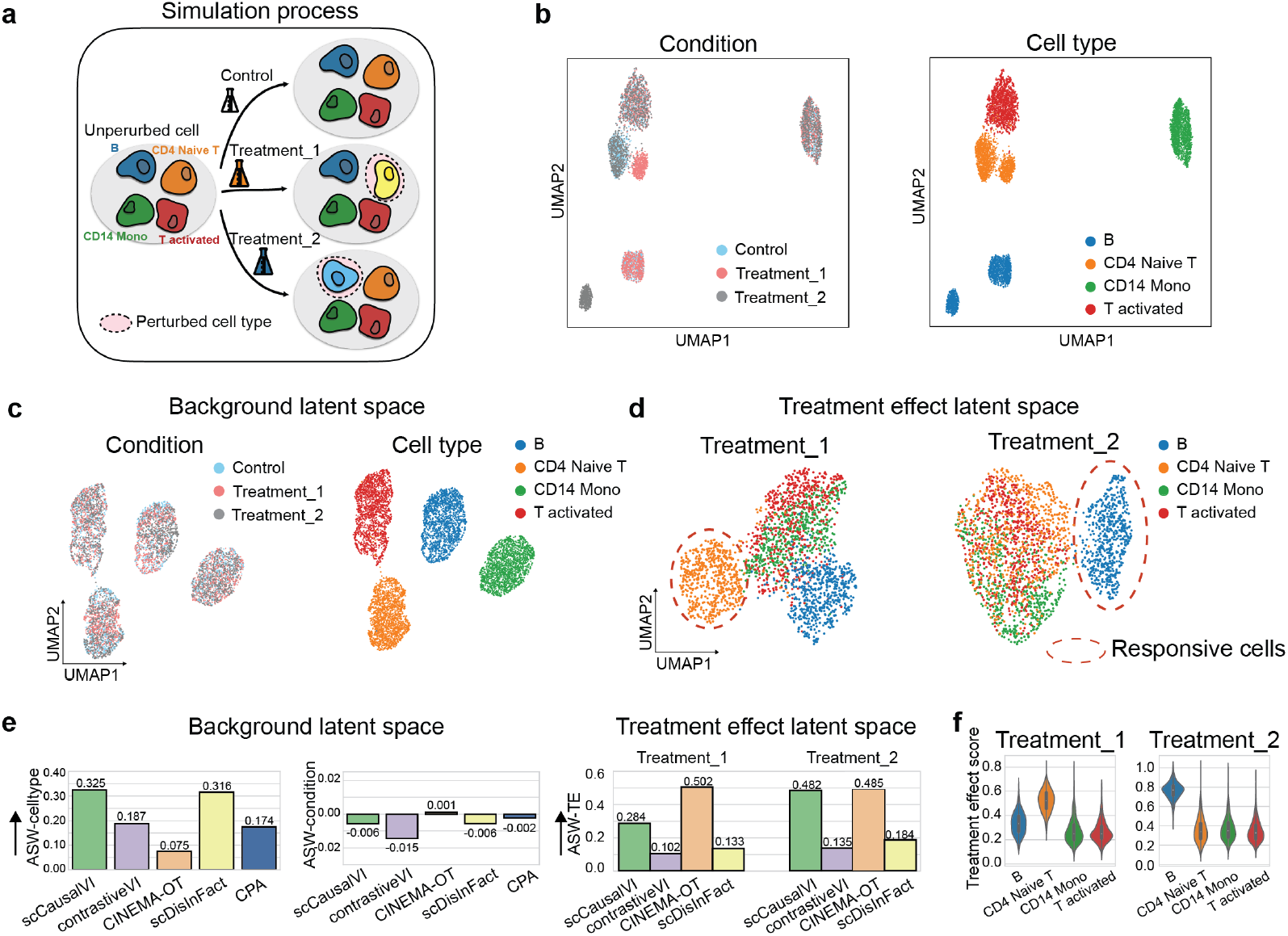
Evaluation of scCausalVI against baseline models on simulated perturbational data. **a**, Schematic of synthetic perturbational data simulation. **b**, UMAP visualization of simulated data labeled by condition and cell type. **c**, UMAP visualization of background latent factors colored by condition and cell type. **d**, UMAP visualization of treatment effect latent factors for two distinct perturbational conditions. Red dashed circles denote responsive populations. **e**, Bar plots showing Average Silhouette Width (ASW)-based metrics for different models and conditions. Left and middle: ASW computed on background latent factors using cell type and condition labels, respectively. Right: ASW computed on treatment effect latent factors. **f**, Distribution of treatment effect scores across cell types.

Next, we analyzed the latent representations by visualization and Average Silhouette Width (ASW)-based metrics to compare performance against baselines in preserving background variation and identifying cell-state-specific treatment effects. In the background latent space, most methods successfully grouped cells by cell type labels and achieved high mixing across conditions, aligning with the ground truth (Fig. 2c, Supplementary Fig. 2). However, CINEMA-OT failed to cohesively group CD4 Naive T cells—which were perturbed in the first treatment condition—in the background latent space, suggesting incomplete isolation of treatment effects from baseline cell states (Supplementary Fig. 2c). The treatment effect latent space revealed more pronounced differences among methods. Only scCausalVI and CINEMA-OT effectively isolated perturbed and unperturbed cells, capturing cell-state-specific treatment effects (Fig. 2d, Supplementary Fig. 2c). In contrast, other baseline methods were hindered by uniform confusion of treatment effects or mixing treatment effects with cell type discrimination (Supplementary Fig. 2a, b, d). The ASW-based metrics demonstrated that scCausalVI outperformed other methods in preserving cell type identity and achieved comparable performance to CINEMA-OT in treatment effect identification (Fig. 2e).

The treatment effect score, a feature only available in scCausalVI, effectively differentiated responsive and non-responsive cell types, with affected populations consistently exhibiting higher scores (Fig. 2f). This quantitative measure, available through our model’s explicit treatment effect latent space, provides a robust means of assessing cellular responses to perturbations at single-cell resolution. To validate the importance of modeling cell-state-specific responses, we performed an ablation study by removing the SENet attention mechanism from scCausalVI. While the model without SENet still effectively captured cell type variations and showed good mixing across conditions in the background latent space, it failed to uncover heterogeneous treatment response patterns (Supplementary Fig. 3). This demonstrates that the SENet attention mechanism is crucial for capturing differential cellular responses to perturbations at single-cell resolution.

### 2.3 scCausalVI outperformed baseline methods in disentangling IFN-*β* responses

We further evaluated scCausalVI and other competing disentanglement methods on a real-world scRNA-seq dataset of IFN-*β*-stimulated PBMC [33]. IFN-*β* stimulation induces widespread transcriptomic changes, observable as shifts in low-dimensional embeddings [33], and cell-type specific responses have been documented through interferon signatures and regulatory pathways [33]. scCausalVI demonstrated outstanding performance in both preserving inherent cellular states in background latent space (Fig. 3a, Supplementary Fig. 4) and identifying differential response (upper-left panel of Fig. 3b). In contrast, the baseline methods were either hindered by cell type identification in background latent space (CINEMA-OT, upper-right panel of Fig. 3b), or failed to discriminate differential cellular response to stimulation (contrastiveVI and scDisInFact, lower panels of Fig. 3b).

**Figure 3:**
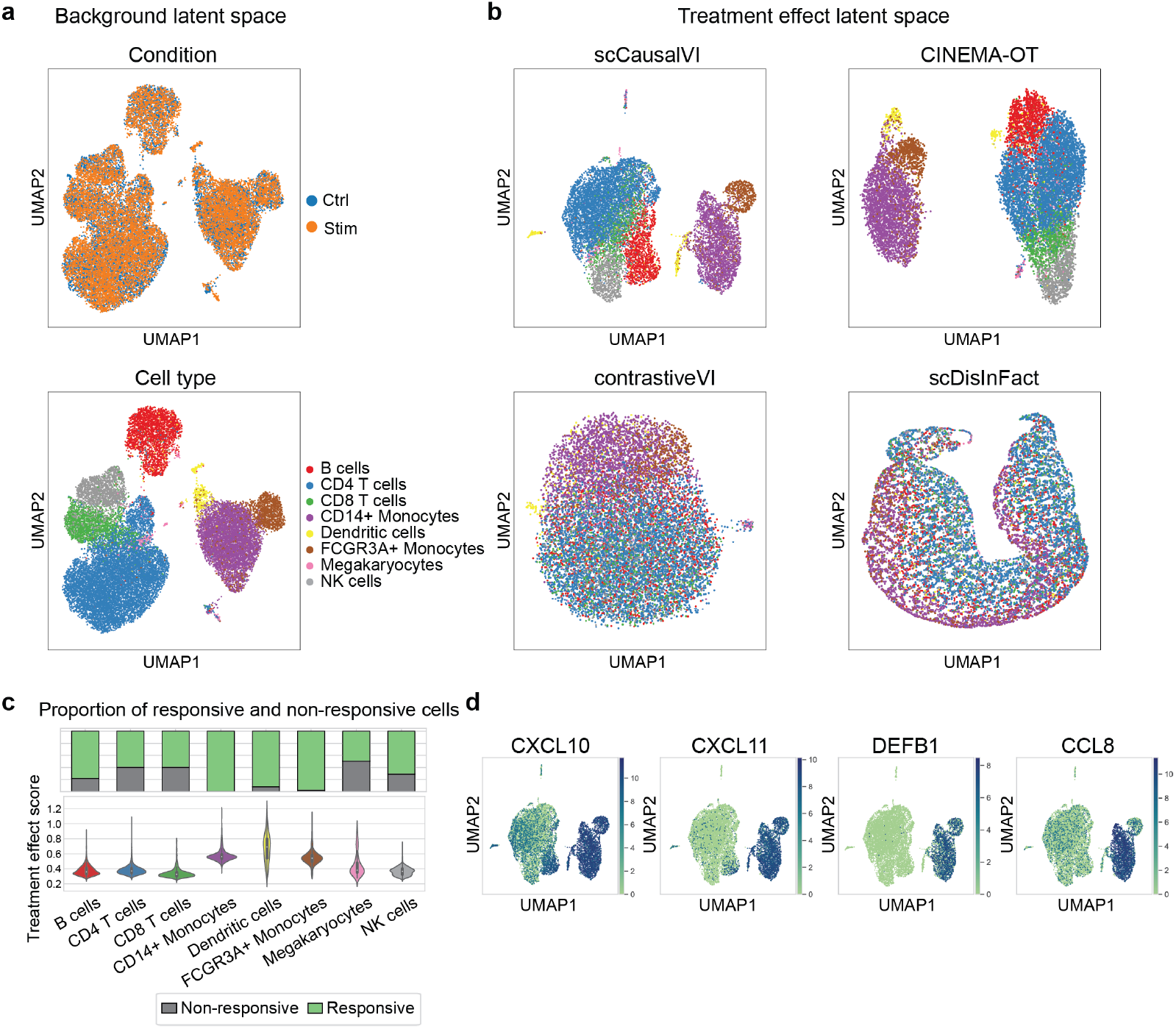
scCausalVI outperformed baseline models in disentangling inherent cellular heterogeneity and differential treatment effects on IFN-*β* data. **a**, UMAP visualization of background latent factors labeled by condition and cell type labels. **b**, UMAP visualization of treatment effect latent factors colored by cell type label by baseline methods. **c**, Distribution of treatment effect scores via violin plots along with the proportion of responsive cells across cell types by bar plots. **d**, UMAP visualization of marker gene expressions by treatment effect latent factors.

Previous studies [33] conducted comprehensive analyses of immune cell responses to IFN-*β*, revealing distinct levels of gene induction across cell types. Specifically, they indicated that monocytes (CD14^+^ and FCGR3A^+^ subsets) and dendritic cells exhibited the strongest upregulation of interferon-stimulated genes in response to IFN-*β*. B cells and T cells still showed evidence of IFN-*β*–driven gene expression changes, but typically these were less pronounced than those observed in the myeloid compartment. By scCausalVI, we quantified the treatment effect size across cell types, along with the proportion of responsive cells in each cell type identified by cross-condition perturbation, which aligns well with the previous findings (Fig. 3c). We further validated the clustering pattern of treatment effect latent factors by examining cell-type-specific responsive markers [7]. These markers displayed distinct expression patterns consistent with known biological responses (Fig. 3d), indicating that scCausalVI effectively captured biologically relevant perturbation response. The consistency between our in silico results and experimental observations underscores the power and reliability of scCausalVI in capturing complex, cell-type-specific responses to cytokine stimulation, demonstrating its potential utility for accurately modeling and interpreting single-cell perturbational data.

### 2.4 scCausalVI accurately performed cross-condition in silico prediction and robustly generalized to unseen cell states

The destructive nature of sequencing methods and population shifts due to perturbations create challenges in the systematic comparison of cellular states across conditions [34]. We evaluated scCausalVI in out-of-distribution (OOD) scenarios, where the model predicts cellular responses for previously unseen cell states, through three key aspects: cross-condition in silico prediction accuracy, generalization to unseen cell types, and robustness to dataset imbalances.

We first benchmarked scCausalVI against the generative baselines with the IFN-*β* stimulated PBMC dataset [33]. scCausalVI achieved superior or comparable cross-condition prediction accuracy (Fig. 4a-c, Supplementary Fig. 5). Moreover, scCausalVI effectively captured significant transcriptomic shifts in marker gene expressions, aligning closely with treated observations, whereas baseline models struggled to accurately predict these changes (Fig. 4d).

**Figure 4:**
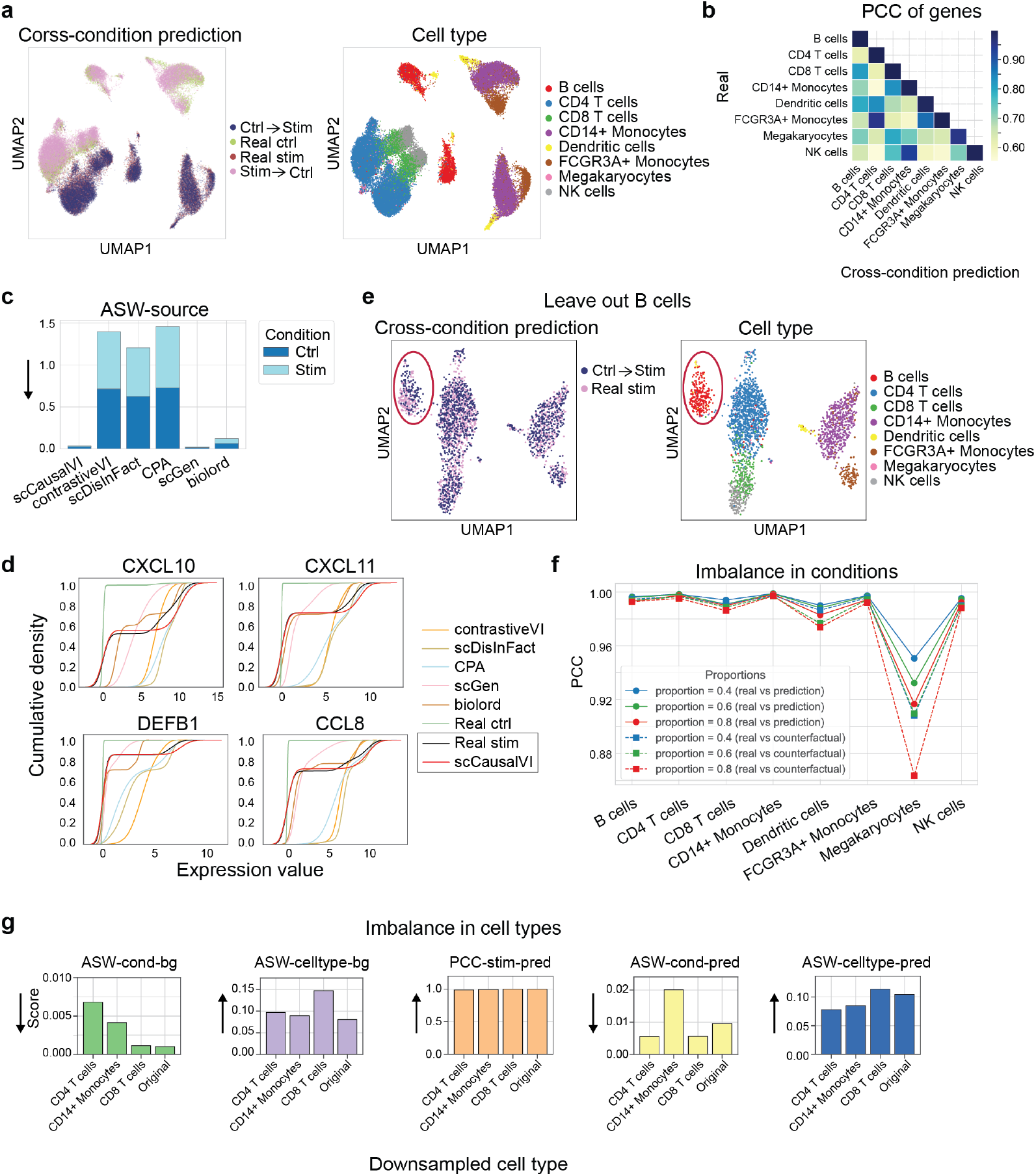
scCausalVI outperformed baseline methods in cross-condition prediction. **a**, UMAP visualization of real and cross-condition predicted data colored by data source and cell type label. “source *→* target” denotes predictions of source cell states under target condition. **b**, Heatmap of PCC of genes between real and cross-condition predicted stimulated data. PCC, Pearson correlation coefficient. **c**, Stacked bar plots representing ASW-based metrics for baselines, assessing the alignment of real and cross-condition predicted data of control and stimulated conditions. **d**, Marginal distributions of marker gene expressions comparing real and cross-condition predictions of stimulated condition by baselines. **e**, UMAP visualization comparing real and cross-condition predictions of stimulated cohort in test data with B cells left out during model training. **f**, PCC between factual predicted (solid lines) or cross-condition predicted (dashed lines) and real gene expression profiles across cell types under various degrees of condition imbalance. **g**, ASW-based and PCC metrics comparing model performance between original data and imbalanced cell type settings, where each cell type in the stimulated condition was downsampled to 10% of its original abundance. The arrows indicate the direction of better performance.

In the OOD settings, cells were randomly split into training (90%) and test (10%) for model training and validation, respectively, with one cell type excluded in training. scCausalVI consistently outperformed other methods in cross-condition prediction (Fig. 4e, Supplementary Fig. 6-12), demonstrating its robust generalization to unseen cell types.

Additionally, scCausalVI demonstrated robust performance under dataset imbalance conditions, encompassing both condition data proportion disparities and alterations in cell type distributions (Fig. 4f–g). In the first scenario, we randomly downsampled the stimulated cell population to 80%, 60%, and 40% of its original size while maintaining the control group unchanged. This created pre-specified imbalance ratios between stimulated and control conditions while preserving their original cell-type compositions. For each level of downsampling, we trained the scCausalVI model on the unbalanced train data, and evaluated its factual prediction and cross-condition prediction accuracy of the stimulated cells from test data. scCausalVI exhibited high robustness, as evidenced by strong Pearson correlation coefficient (PCC) across cell types (Fig. 4f) and consistent UMAP visualizations (Supplementary Fig. 13).

In the second scenario, we assessed scCausalVI’s robustness to imbalances in cell type distributions by selectively downsampling three predominant cell types—CD4 T cells, CD8 T cells, and CD14+ monocytes—to 10% of their original abundance within the treated condition. This significant reduction introduced a pronounced imbalance of these cell types between control and stimulated conditions, challenging the downstream analyses after integration [35]. Despite this extreme imbalance, scCausalVI maintained its performance in disentangling latent factors and ensuring accurate factual prediction and cross-condition prediction (Fig. 4g, Supplementary Fig. 14). Compared to results obtained using the complete original dataset, the downsampled data revealed a distinct separation of B cells from other lymphoid cell types (CD4 T, CD8 T, NK) and megakaryocytes in the treatment effect latent space. This separation likely reflects heterogeneity in cellular responses at a more granular level, as evidenced by the distinct expression patterns of the marker gene *CXCL10* in B cells versus other lymphoid cells (Fig. 3d). This robustness and accuracy demonstrated scCausalVI’s potential as a powerful tool for modeling complex single-cell perturbation data, facilitating deeper insights into cellular heterogeneity and treatment effects.

### 2.5 scCausalVI effectively disentangled treatment effects from batch effects in multi-source data integration

The increasing amounts of data available from different laboratories necessitate robust methods for integrative analysis while preserving biological signals and removing technical variations [36]. We evaluated scCausalVI’s capacity to distinguish between genuine biological treatment effects and technical batch effects using two independent PBMC scRNA-seq datasets from COVID-19 studies. These datasets, generated by Meyer et al. [37] and Blish et al. [38], respectively, comprised samples from both healthy donors and COVID-19 patients. To maintain compliance with randomized experimentation assumptions, we focused our analysis on eight cell types that were present in both conditions and both datasets, creating a combined dataset spanning two experimental conditions (healthy and COVID-19) and two distinct batch sources (Blish and Meyer).

Initial analysis revealed substantial batch effects in the original data space, manifesting as a clear separation between the two datasets (Supplementary Fig. 15a). While conventional batch correction with Harmony [39] successfully aligned the datasets, it preserved the underlying treatment effects in the corrected embeddings (Supplementary Fig. 15b). scCausalVI, through explicit encoding of batch indices, achieved comprehensive disentanglement of cell states, treatment effects, and batch effects. The background latent space revealed well-preserved inherent cell heterogeneity, with cells clustering by annotated cell type labels and demonstrating thorough mixing across both batches and conditions (Fig. 5a). We quantified the mixing across conditions and batches by entropy score and cell type identity by silhouette width. scCausalVI’s performance was superior or comparable to Harmony-corrected embedding (Fig. 5b). Furthermore, the treatment effect latent space exhibited complete batch-independent clustering of treated cells (Fig. 5c), confirming the effective isolation of treatment effects from batch effects.

**Figure 5:**
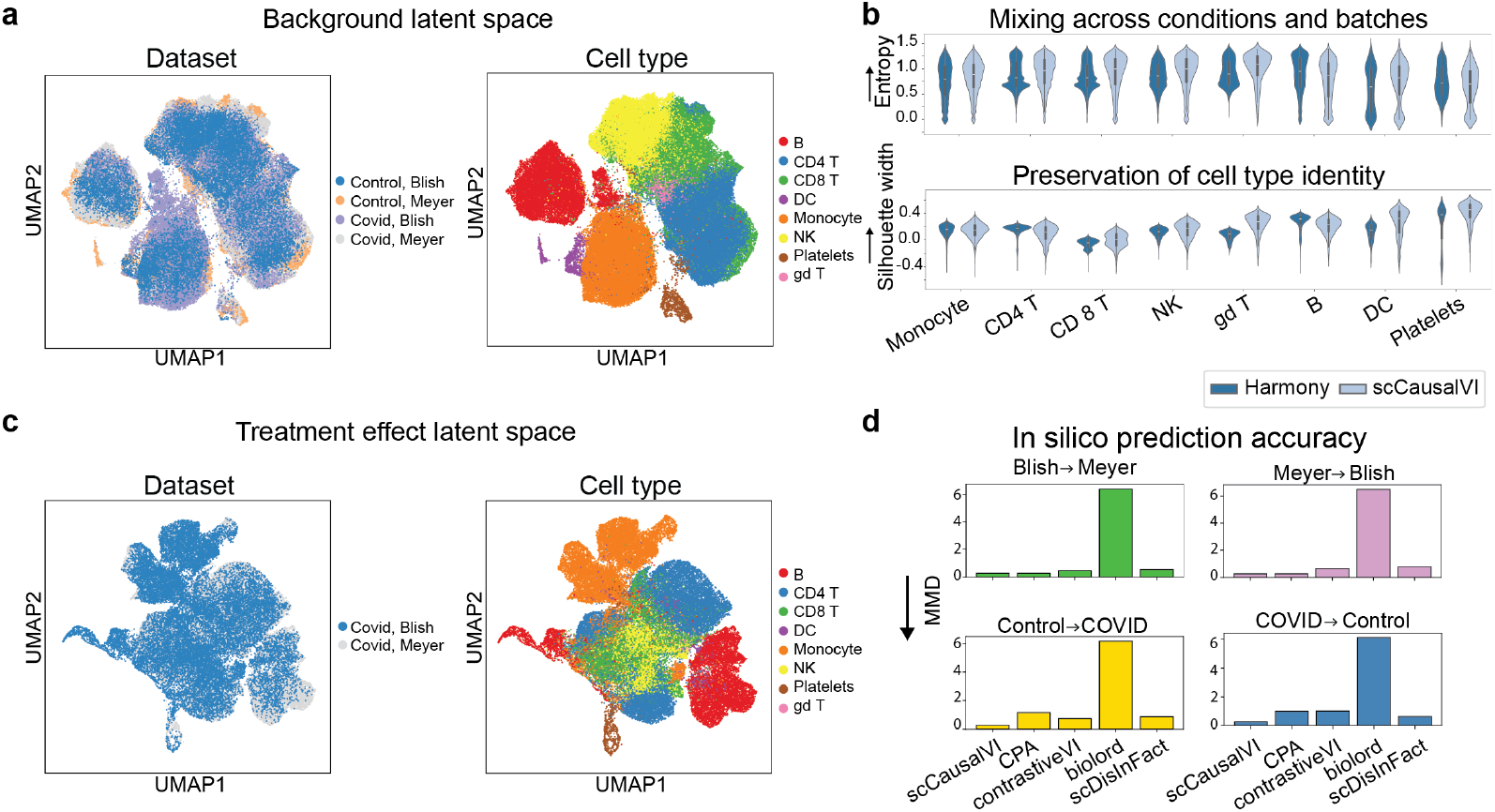
scCausalVI analyses of multi-batch COVID-19 PBMC datasets. **a**, UMAP visualization of the background latent factors colored by dataset source and cell type labels. **b**, Entropy scores for mixing across conditions and batches (top), and silhouette width scores by cell type labels (bottom) with Harmony embedding and background latent factors of scCausalVI, respectively. **c**, UMAP visualization of treatment effect latent space colored by dataset source and cell type labels. **d**, MMD between distributions of in silico predictions and target observed populations calculated on the top 50 principal components of their joint PCA, compared across methods for both cross-batch and cross-condition predictions. A lower MMD indicates higher prediction accuracy. MMD, maximum mean discrepancy.

To quantitatively assess scCausalVI’s ability to distinguish between batch effects and treatment effects, we performed in silico predictions by independently manipulating batch indices and treatment assignments. For cells from any condition or batch, we generated in silico predictions under altered batch or condition settings. For each pair of predicted and target observed populations, we performed principal component analysis (PCA) on the concatenated data and measured the maximum mean discrepancy (MMD) on the top 50 principal components between predictions and observations for each cell type. scCausalVI displayed remarkable accuracy and outperformed other generative baselines (Fig. 5d), further validating its performance in disentangling treatment effects and batch effects.

The robustness of scCausalVI was further validated by the consistency between parallel integrative and separate analyses of the two batches. Both analytical approaches showed substantial agreement in responsive cell identification patterns (Supplementary Fig. 16a,b), with hypergeometric tests confirming significant overlap (*p <* 0.001) and observed-to-expected ratios of 3.96 and 1.84 for Meyer and Blish datasets, respectively. When comparing responsive and non-responsive cells, both analytical approaches identified similar sets of differentially expressed genes from responsive samples, with Jaccard similarity scores of 0.76 and 0.78 between the top 200 genes from the integrative and separate analyses in the Meyer and Blish datasets, respectively (Supplementary Fig. 16c). These findings support scCausalVI’s robustness and reliability in separating treatment-induced biological signals from technical variations across different analytical strategies in multi-source single-cell datasets.

### 2.6 Negative control validation revealed robust batch effect handling in scCausalVI

To evaluate scCausalVI’s ability to distinguish technical batch effects from genuine biological signals, we conducted a negative control experiment using control samples from two independent batches (Blish and Meyer) (Supplementary Fig. 17a,b). This experimental design, treating technical batches as pseudo-conditions, allowed us to assess whether the model would erroneously interpret batch-specific variations as treatment effects.

We performed analysis by setting dataset of two (pseudo-) conditions and two batches. The background latent embeddings generated by scCausalVI effectively served as batch-corrected representations, demonstrating the successful integration of the two batches while maintaining the underlying biological structure of distinct cell populations (Supplementary Fig. 17c). Comparative analysis with Harmony revealed that scCausalVI achieved comparable or superior performance in both batch mixing and cell type identity preservation, as quantified by entropy scores and silhouette width metrics, respectively (Supplementary Fig. 17d).

In this negative control context, scCausalVI demonstrated high specificity, correctly classifying the majority of cells as non-responsive. However, we observed elevated false positive rates in specific cell populations (Supplementary Fig. 17e). Dendritic cells exhibited slightly higher false positive rates, attributable to their inherent transcriptional heterogeneity evidenced by distinct subpopulations in the original data (Supplementary Fig. 17a,f). Similarly, platelets showed increased false positives, likely due to their relative scarcity in the dataset and unique biological characteristics as anucleate cells with minimal transcriptional activity, rendering them particularly susceptible to technical variations.

To further validate scCausalVI’s in silico perturbation capabilities, we performed systematic interventions on condition and batch indices independently. Given the absence of true treatment effects between control datasets, batch index manipulation should yield predictions aligned with target batch distributions, while condition index alterations should maintain concordance with source data. Quantitative assessment using MMD on the top 50 principal components demonstrated superior alignment between predicted and target populations compared to Harmony-based integration (Supplementary Fig. 18a). UMAP visualization corroborated these findings, showing consistent alignment between in silico perturbed data and corresponding target populations across experimental conditions (Supplementary Fig. 18b). These comprehensive negative control analyses demonstrate scCausalVI’s robust capability to effectively distinguish and handle batch effects without erroneously attributing technical variations to treatment effects, thereby validating its utility for accurate interpretation of multi-batch single-cell datasets.

### 2.7 scCausalVI discriminated responsive and non-responsive respiratory epithelial cells to COVID-19 by in silico perturbation

Traditional analysis of perturbation experiments is limited by relying solely on experimental condition labels, where all cells in the treated condition are assumed to be uniformly affected. However, biological variability in response to treatment—influenced by factors including off-target effects, cell cycle states, metabolic conditions, and the microenvironment—can result in differential responses even among cells of the same type. This binary classification based on experimental conditions can obscure significant differences between responsive and control populations [40]. To address this issue, we developed statistical methods within scCausalVI to distinguish between responsive and non-responsive cells in treated groups. In the following experiments, we did not observe significant batch effects, hence no batch index was included in the model.

We first validated scCausalVI’s ability to identify treatment-responsive cells within the treated cohort using a COVID-19 dataset of respiratory epithelial cells from both healthy donors and patients [41]. Responsive cells are defined as those exhibiting measurable changes in response to treatment, either through direct effects on primary treatment targets or indirectly via secondary effects or intercellular interactions. In this dataset, we considered the cells with detectable viral transcripts as responsive. scCausalVI classified cells as responsive or non-responsive by comparing observed profiles with both factual and cross-condition predictions, achieving 98% precision for non-responsive and 11% precision for responsive cell identification (Supplementary Fig. 19a,b). In contrast, no cells were classified as responsive by all the other baselines following the same in silico perturbation-based procedure.

We investigated cells where scCausalVI’s responsiveness predictions diverged from viral transcript detection labels. Analysis of COVID-19-associated markers (interferon-stimulated genes and SARS-CoV-2 receptor *ACE2*) [42] revealed that cells predicted as responsive by scCausalVI, despite lacking viral transcripts, exhibited elevated expression of these markers (Supplementary Fig. 19c). Conversely, cells containing viral transcripts but classified as non-responsive by scCausalVI showed expression patterns similar to true non-responsive cells. These findings suggest our model could potentially improve the accuracy of identifying responsive cells beyond viral transcript detection alone, offering a more comprehensive identification of cellular responses.

### 2.8 Cross-condition in silico prediction facilitated heterogeneity of susceptible monocytes to COVID-19

To further support and expand the power and potential of in silico perturbation, we applied scCausalVI to the COVID-19 dataset of PBMC cells from Blish et al. that lacks explicit labels for responsive or non-responsive [37]. We aimed to study the differences between susceptible and resistant cells to COVID-19 in the healthy state, enabling us to identify the putative drivers of disease susceptibility. This approach provides insights that go beyond conventional association studies, offering a more robust foundation for understanding disease mechanisms and developing targeted therapies.

In the latent representation by scCausalVI, we observed a substantial mixing of cells from different conditions, with clustering by annotated cell type labels in the background latent space (Fig. 6a). Notably, significant heterogeneity in treatment effect latent factors persisted within cells of the same type (Fig. 6b-c, Supplementary Fig. 20a). The alignment between the factual and cross-condition predictions with the observed data (Supplementary Fig. 20b-c) further underscored the model’s reliability for downstream analyses. Remarkably, the identified non-responsive cells from COVID-19 patients largely overlapped with cells from healthy donors. In contrast, responsive cells exhibited distinct shifts indicative of disease-related changes (Fig. 6d). Quantitative analysis using *k*-nearest neighbors (*k* = 30) on the top 20 principal components revealed that among disease cells, on average only 14.17% of those proximal to healthy cells were responsive, while 85.83% were non-responsive, significantly deviating from the null distribution obtained through permutation testing (*n* = 500; 51.07% responsive, 48.93% non-responsive, proportional to the overall treated cohort composition). This pattern revealed through scCausalVI’s in silico perturbation, suggested that non-responsive cells maintain a phenotype similar to healthy cells, while responsive cells undergo significant alterations in response to infection.

**Figure 6:**
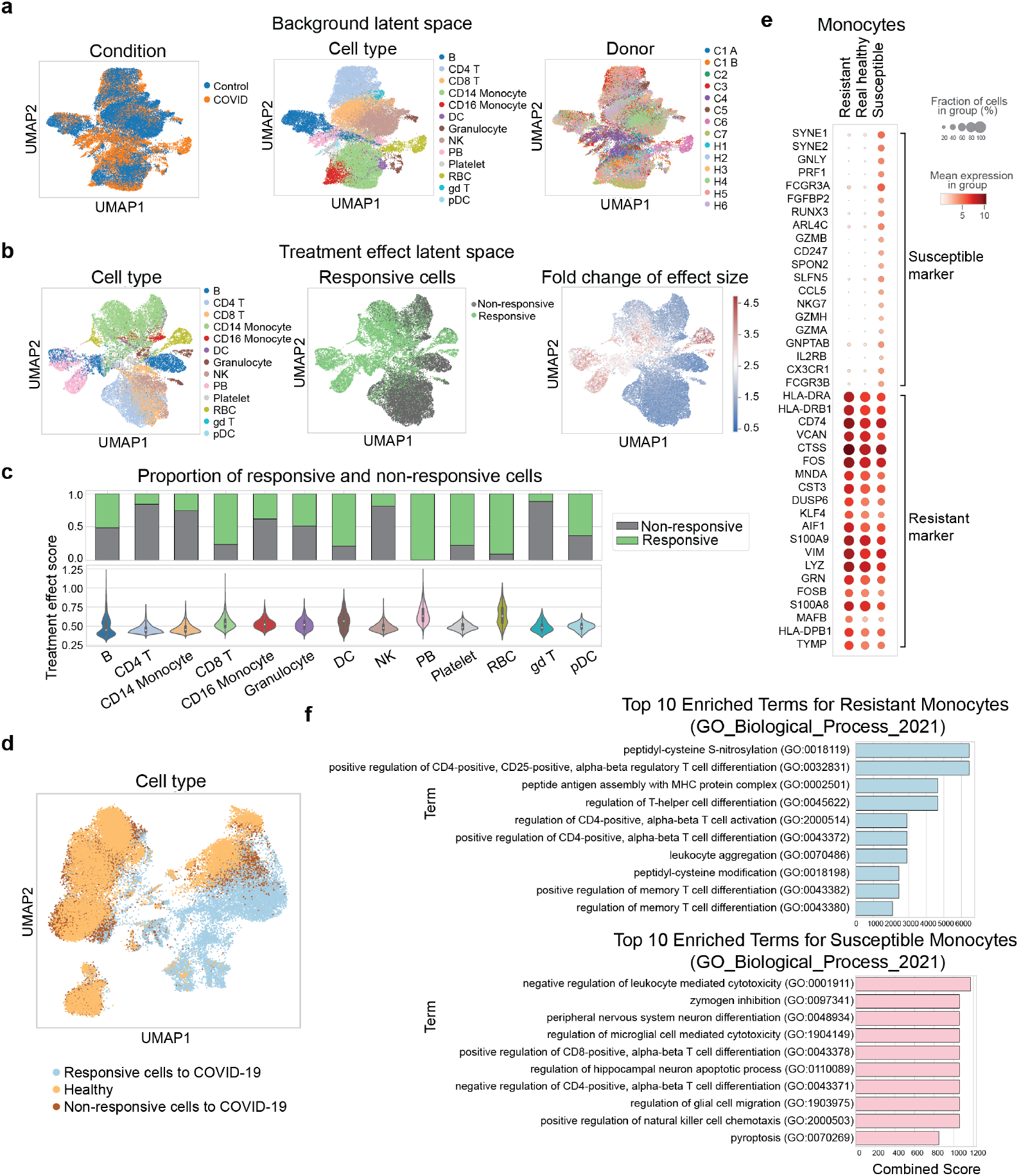
scCausalVI facilitated characteristics of susceptible and resistant PBMC cells to COVID-19. **a**, UMAP visualizations of the background latent factors, colored by condition, cell type label provided by [37], and donor. **b**, UMAP visualizations of treatment effect latent factors, colored by cell type label, predicted responsive cells, and fold change of effect size. **c**, Distribution of treatment effect scores across cell types via violin plots and proportion of responsive cells by bar plots. **d**, UMAP visualization of the entire dataset, colored by healthy versus COVID-19 labels and indicating the identification of responsive versus non-responsive cells within the infected cohort. **e**, Dot plot for differential gene expression in real healthy cells, predicted susceptible and resistant cells in monocytes. **f**, Top 10 enriched GO terms for predicted resistant and susceptible monocyte cells.

We focused on monocytes, which play a crucial role in immune regulation and show significant variability in response to SARS-CoV-2 infection. Using cross-condition in silico prediction, we generated expression profiles for responsive and non-responsive monocytes under healthy conditions, computationally identifying susceptible and resistant cell states. Among the two populations, the differential expression patterns and phenotypes were identified with Wilcoxon rank-sum test (Fig. 6e, Supplementary Fig. 20d). Specifically, we observed upregulation of *FCGR3A* and *FCGR3B* in susceptible cells, which were previously described as CD16+ intermediate or non-classical monocytes, and reported as depleted in COVID-19 patients [43, 44, 45]. This finding helps explain how elevated baseline inflammatory potential might predispose cells to more aggressive inflammatory responses upon viral challenge, potentially increasing susceptibility to severe outcomes in infections like COVID-19. Conversely, resistant cells exhibited high expression of *HLA-DRA* and *HLA-DRB1* [46, 43]. It has been shown that elevated expression of these MHC class II genes in monocytes is associated with favorable prognosis in COVID-19 patients, suggesting that maintenance of antigen presentation capacity may be a key factor in cellular resistance to severe COVID-19 outcomes.

Gene Ontology (GO) term analysis of differentially expressed genes further illuminated the molecular basis of cellular resistance and susceptibility to COVID-19 (Fig. 6f). In resistant cells, we observed enrichment of terms related to peptidyl-cysteine S-nitrosylation and modification, aligning with recent findings on potential therapeutic targets for inhibiting SARS-CoV-2 infection [47]. Enrichment in peptide antigen assembly with MHC protein complex and regulation of CD4-positive, alpha-beta T cell activation can help in mounting an effective immune response, potentially reducing the severity of the disease and aiding in the protection of uninfected cells [48, 49]. Conversely, susceptible cells showed enrichment in terms associated with negative regulation of leukocyte-mediated cytotoxicity which is reasonable since impaired cytotoxic responses can lead to ineffective viral clearance [50]. Notably, the enrichment of pyroptosis-related genes in susceptible cells aligns with recent research identifying inflammasome activation and cellular pyroptosis as promising targets for treating severe COVID-19 [51].

## 3 Discussion

To tackle the challenge of disentangling treatment effects from inherent cellular heterogeneity in perturbational scRNA-seq data, we developed scCausalVI, a causality-aware generative model that separates causally related variations at single-cell resolution. scCausalVI utilizes causal disentanglement to distinguish cellular heterogeneity from treatment-specific effects and employs attention mechanisms to adaptively capture differential response patterns. The causality-aware design of scCausalVI provides mechanistic insights into cellular responses and an interpretable latent representation of the dynamic processes.

A key feature of scCausalVI is cross-condition prediction, which enables in silico perturbations to predict gene expression under hypothetical conditions, offering the ability to direct comparison of cellular states across conditions at the single-cell level. scCausalVI demonstrated superior performance in capturing differential treatment responses, out-of-distribution generalization, and robustness under data imbalances compared to state-of-the-art methods. The robustness makes it well-suited for precision medicine applications, accurately characterizing differential responses even for underrepresented cell types, and distinguishing between responsive and non-responsive cells.

Notably, a significant advancement of scCausalVI lies in its ability to simultaneously disentangle batch effects, treatment effects, and cellular baseline states in multi-source single-cell perturbational data integration. Through explicit encoding of batch indices, scCausalVI achieves comprehensive separation of these intertwined variations, validated by integrative analyses of multi-batch data and negative control experiments. Furthermore, in silico prediction through independent manipulation of batch and condition indices demonstrated precise alignment with target distributions, outperforming generative baseline models. These capabilities are particularly valuable for modern single-cell studies, where integrative analyses of data from multiple sources are increasingly common and the distinction between technical artifacts and biological signals is crucial for accurate interpretation [36].

Despite the strengths of scCausalVI, there are opportunities to enhance the model’s effectiveness. Although regularization techniques such as MMD and *L*2-norm constraints are currently used, future work could explore employing identifiable models for causal disentanglement by integrating them with deep latent-variable models [52, 53]. Additionally, extending scCausalVI to better accommodate largescale perturbation experiments presents a compelling research avenue. The current architecture involves separate encoders for each condition, which needs to be further optimized for scalability when dealing with multiple perturbations, as seen in genome-wide CRISPR studies [4, 54]. One potential strategy is to use a foundation model to encode data from different conditions, significantly enhancing scalability and making the model applicable to large-scale perturbation experiments [29, 55].

In summary, scCausalVI provides a robust and interpretable framework for disentangling cellular heterogeneity from cell-state-specific treatment effects at the single-cell level. Its ability to perform in silico perturbation opens new avenues for understanding the causal mechanisms underlying cellular responses and offers promising applications in the identification of therapeutic targets and the development of personalized treatment strategies. By addressing the challenges of causal disentanglement and predictive generalization, scCausalVI represents a significant advancement in single-cell analysis, pushing the boundaries of what can be inferred from perturbational experiments.

## 4 Methods

scCausalVI is an unsupervised causality-aware generative model for disentangling and modeling causal mechanisms underlying perturbational scRNA-seq data.

### 4.1 Framework of scCausalVI

#### 4.1.1 Theoretical background

We consider case-control scRNA-seq experiments in which cells are randomly assigned to either control (untreated) or one of multiple treatment conditions. Our randomized experimental design ensures two critical causal inference assumptions. First, treatment assignment is independent of inherent cellular states (ignorability), eliminating bias from pre-existing cellular differences in causal effect estimation. Second, the Stable Unit Treatment Value Assumption (SUTVA) is satisfied through the physical separation of cells across treatment conditions, preventing interference between treatment groups. This experimental setup provides a rigorous foundation for unbiased causal inference in perturbational singlecell studies.

Based on these assumptions, we incorporate Structural Causal Models (SCMs) with variational inference to model causal mechanisms with unobserved latent factors. Given cells’ baseline states in the untreated group *z*, treatment-induced cell-state-specific response *e* under treatment *t*, and covariates such as batch index *c* if applicable, the SCM learns the causal mechanism generating observed values *x* from these components:

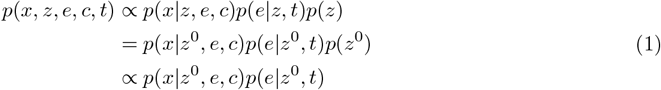

Here, *z*^0^ represents the background latent factors from the control group. Under our randomized experimental design, all background latent factors *z* from treatment conditions follow the same distribution as *z*^0^, reflecting the independence between treatment assignment and inherent cellular states. The model captures how treatment effects *e* are generated conditional on baseline states *z*, while the observed expression profile *x* depends on both the baseline state and treatment response.

#### 4.1.2 Generative process of scCausalVI

To account for the possible uncertainty, we analyze the sequenced count data by a probabilistic model. Without loss of generality, the control group is set as the first condition, namely *t* = 0 for control data 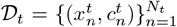 and *t >* 0 for cells upon perturbation or treatment 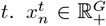 denotes the single-cell profile of *G* genes from *t*th condition, and 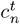 is the categorical covariate, e.g., the experimental batch index. Each perturbed expression value 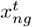 from *t*th perturbational condition (*t >* 0) is drawn independently through the following process:

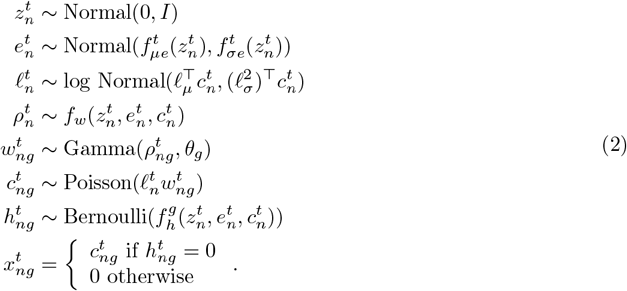

In this process, 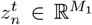 and 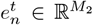 refer to the two sets of latent variables underlying the condition-specific scRNA-seq data, which we designate as the background latent factors and the treatment effect latent factors, respectively. The latent variable 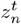 captures the intrinsic cellular heterogeneities representing the unperturbed, healthy, or control states of cells, which are shared across both control and various treatment conditions. In essence, 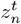 encodes the inherent cellular states as they exist in the absence of any treatment or perturbation. The latent factor 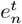 represents the treatment effects specific to each condition, capturing how perturbations or treatments modulate these intrinsic cellular states. For each specific condition *t >* 0, a multilayer perceptron 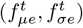 is employed to encode the condition-specific generative process that transforms the unperturbed cell states 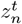 into the corresponding treatment effects 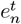 at single-cell resolution. In the control group, cellular heterogeneities are fullycharacterized by 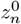, and we set 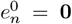 to denote the absence of treatment effects. No generative process 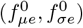 is applied to control data, as there are no perturbations to the model. In summary, the background latent factors *z* capture the underlying heterogeneities of unperturbed states of scRNA-seq data. For treated data (conditions *t >* 0), both 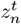 and 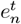 are required to account for biological variations—where 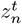 represents the cells’ baseline, unperturbed states, and 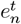 captures the deviations due to treatments or perturbations.

Here, *f*_*w*_ and *f*_*h*_ in the generative process map the latent space and batch annotations to the original gene space, i.e., 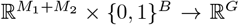. The network *f*_*w*_ is constrained during inference to encode the mean proportion of transcripts expressed across all genes using a softmax activation function in the layer. And network *f*_*h*_ encodes whether a particular entry has dropped out owing to technical effects. *B* denotes the cardinality of the categorical covariate and *c*_*n*_ ∈ {0, 1}^*B*^ represents categorical covariates, such as experimental batches. For each category, 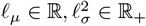 parameterize the prior for the scaling factor on a log scale, and are set to be the empirical mean and variance of the log-library size of each corresponding batch, respectively. The variable 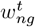 represents the underlying true expression level, 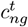 is the raw count, and 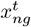 is the final observed expression value after accounting for dropouts via 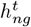. This hierarchical structure induces a Zero-Inflated Negative Binomial (ZINB) distribution for the observed counts 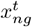, as the Gamma-Poisson mixture marginalizes to a Negative Binomial distribution. The ZINB formulation effectively captures both overdispersions through the Negative Binomial component and excess zeros through the zero-inflation component, characteristic features of single-cell RNA sequencing data.

#### 4.1.3 SENet attention module

To better account for the differential treatment effect and the varying magnitude of each single cell, scCausalVI incorporates an adaptive feature scaling mechanism inspired by Squeeze-and-Excitation Networks (SENet) [30]. We adapt this mechanism for treatment effect estimation in case-control studies to capture the complexities and heterogeneities of cellular responses to treatments.

In the treatment effect latent space, we employ a sub-network to generate scaling parameters for the latent factors. Formally, given the treatment effect latent factors 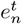, the scaling operation is defined as:

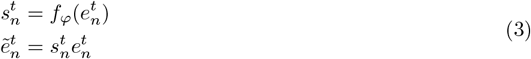

where *f*_*φ*_ is the scaling network comprising a single linear layer parameterized by *φ*, followed by a Softmax activation function. The output 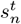 represents the generated scalar scaling parameter. The rescaled treatment effect latent factors 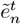 are subsequently utilized in the generative process of treated measurements.

This approach allows the scaling parameter 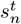 to be interpreted as a measure of the magnitude of treatment effect in a specific cellular context, enabling fine-grained modeling of condition-specific responses at single-cell resolution. Consequently, it serves as a quantitative indicator of differential perturbation responses across various cellular states. The importance of this rescaling mechanism has been rigorously validated through ablation studies, demonstrating its crucial role in accurately capturing cell-state-specific treatment effects.

By implementing this explicit generative process to describe the mechanisms of treatment-based single-cell experiments, our method aims to learn causally disentangled and semantically meaningful representations - specifically, the inherent cellular heterogeneities and cell-state-specific treatment effects. The generative framework enables us to perform in silico perturbation and estimate potential expression profiles of unseen cells at single-cell resolution, thereby extending the utility of our model beyond the observed data.

#### 4.1.4 Inference of scCausalVI

The posterior cannot be obtained directly by using Bayes’s rule because computing the integrals required for 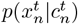 in the denominator is analytically intractable. As in scVI, we instead approximate the posterior distribution using variational inference. For the *n*th observation 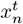 from *t* th treatment, the variational posterior 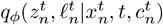 is factorized with the mean-field approximation:

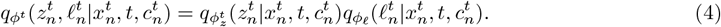

We set the variational posterior 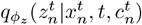 to be Gaussian with a diagonal covariance matrix, with neural network framework 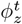 to encode the mean and covariance with input 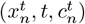 [56]. The posterior distribution of 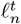 is chosen to be the log-normal with the scalar mean and variance encoded by a neural network 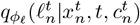. The evidence lower bound is

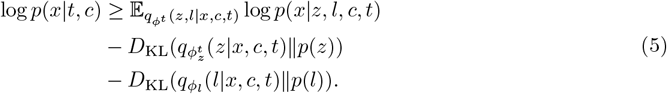

For data sampled from the control group, the treatment effect latent factor 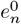 is set to **0**, thus the likelihood *p*(*x* |*z, l, c, t* = 0) can be simplified by integrating out *w*_*ng*_, *h*_*ng*_ and *c*_*ng*_, yielding a probability following a Zero-Inflated Negative Binomial (ZINB) distribution. That is for *t* = 0,

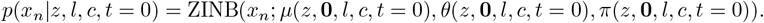

The generative process of perturbational data involves the mediate hidden factor *e*, and the likelihood for data 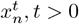 is

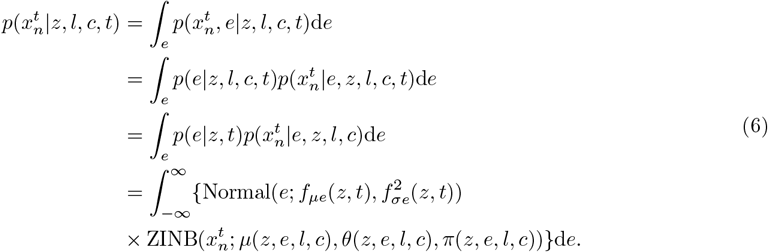

Since the integral is intractable due to the complexity introduced by neural networks in the generative process, we approximate the marginal likelihood using Monte Carlo estimates as follows:

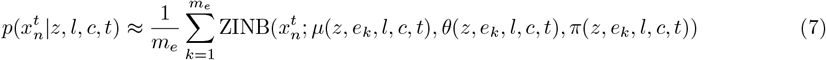

with each *e*_*k*_ sampled from distribution Normal 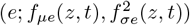.

### 4.2 Loss functions to encourage causal disentanglement and capture cell-to-cell variability in treatment effects

Overall, scCausalVI contains multiple networks tailored for the control group and each of the treated conditions. For data from each condition, a specific encoder is used to infer the background latent factors. Additionally, an extra generative module is included to model the treatment effect latent factors for each treated group, capturing cell-state-specific treatment effects. With the assumption of a randomized case-control design, the background latent factors are expected to follow an identical distribution across both control and treated groups. To enforce this alignment of the background latent factors and promote causal disentanglement of confounding factors from treatment effects, we employ a Maximum Mean Discrepancy (MMD) regularizer [57] on their distributions, encouraging the causal disentanglement of confounding factors from treatment effects [58]. Specifically, we use the background latent factors 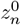 from the control data as a reference to align those from treated conditions:

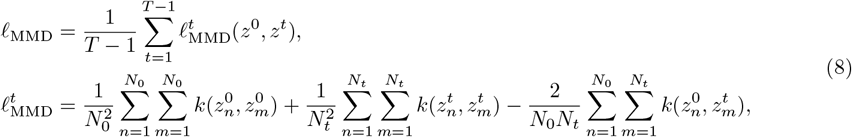

where *N*_*t*_ is the number of cells in condition *t* and *T* is the number of experimental conditions including the control group. *k*(*·, ·*) denotes a Gaussian radial basis function kernel defined as:

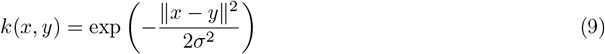

where *σ* is the kernel bandwidth parameter and ∥*x* − *y*∥ ^2^ is the squared Euclidean distance between points in the latent space.

Despite the efforts to align confounding factors, the inherent non-identifiability of neural networks poses challenges in achieving complete causal disentanglement between intrinsic cellular heterogeneity and cell-state-specific treatment effects. The treatment effect latent space may inadvertently inherit variations from the background latent space, potentially confounding treatment effects with inherent cellular characteristics. To mitigate this issue and enhance the fidelity of the revealed treatment effects, we introduce a regularization term that constrains the magnitude of the treatment effect latent factors. This strategy aims to prevent information leakage from the background latent space into the treatment effect representation. Specifically, we implement an *L*2-norm regularization on the treatment effect latent factors:

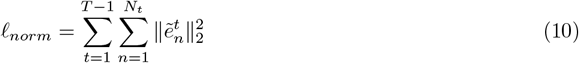

where 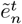 represents the treatment effect latent factors. This regularization term encourages the model to capture only the essential treatment-induced variations, thereby promoting a more focused and interpretable representation of treatment effect latent factors.

Therefore, the final loss function of our algorithm is defined as

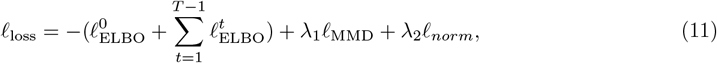

where

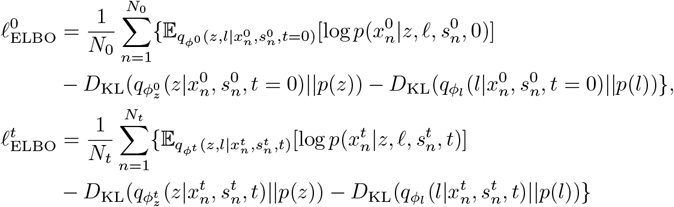

with the penalty parameter *λ*_1_ *>* 0, *λ*_2_ *>* 0 for encouraging causal disentanglement and capturing differential treatment effects.

### 4.3 Optimization details

The framework of scCausalVI comprises condition-specific modules and shared modules depending on whether the module forwards data from one specific condition or all conditions. The condition-specific modules are trained separately with data from their respective conditions, while the shared modules are trained with data from all conditions.

The implementation of scCausalVI is based on the scvi-tools (version 0.16.1) python package, utilizing the Encoder and DecoderSCVI modules. For all datasets used in this research, we adopted the default parameter settings of scCausalVI. The Encoder network consists of two fully connected layers with ReLU activation functions, processing the expression values concatenated with the one-hot encoding of covariates (e.g., batches). Each hidden layer contains 128 neurons, and the background latent factors are set to follow a 10-dimensional Gaussian distribution. The generative network, which connects the background latent space to the treatment effect space, is composed of two hidden layers with ReLU activation. The treatment effect latent factors are also set to a 10-dimensional embedding. The DecoderSCVI mirrors the encoder network, containing 2 layers with ReLU activation and 128 neurons in each hidden layer. The input of the decoder is the concatenated background and treatment effect latent factors, as well as the one-hot encoding of covariates if available, and the output is the parameters of ZINB distribution.

Parameter *λ*_1_ of MMD loss, which enforces the alignment of unperturbed cell states in background latent space, demonstrates robustness across values ranging from 1 to 10. We set *λ*_1_ = 10 as the default value, which consistently achieves effective alignment of background latent factors across all our analyses. Parameter *λ*_2_ of *L*2 norm regularization on treatment effect latent factors penalizes the expressiveness of cell-type variance in the treatment effect space. Lower values of *λ*_2_ permit greater variability in treatment effect latent factors, while higher values compress less significant variations. Through empirical testing, we found *λ*_2_ performs optimally in the range of 0.2 to 0.4, with 0.3 recommended as the default value due to its balanced regulation of treatment effect representations. The model is trained using the Adam optimizer with an initial learning rate of 0.001. Early stopping is employed to prevent overfitting, with a patience of 10 epochs and a maximum of 500 training epochs.

### 4.4 Downstream analysis of scCausalVI

#### 4.4.1 Analysis of the disentangled latent factors

##### Clustering and visualization

By disentangling the background latent factors and the treatment effect latent factors, our model supports standard scRNA-seq analyses. Firstly, clustering in the background latent space allows for the identification of cellular compositions in their unperturbed or baseline states. This analysis reveals inherent cellular heterogeneity without the confounding influence of treatments or perturbations. Secondly, clustering within the treatment effect latent space enables the detection of differential cellular responses to treatments. This approach uncovers subpopulations of cells that exhibit distinct responses, highlighting cell-state-specific treatment effects. Visualization of these latent spaces using dimensionality reduction techniques (e.g., t-SNE, UMAP) provides intuitive representations of how cells differ intrinsically and in their responses to treatments.

##### Quantitative metric of treatment effect size

To quantitatively assess the magnitude of treatment effects at the single-cell level, we compute the *L*2-norm of the treatment effect latent factors. This metric serves as a measure of treatment effect size in the latent space, allowing for the identification of cells with varying degrees of responsiveness.

#### 4.4.2 Cross-condition in silico prediction by causal generative model

##### In silico prediction of unseen cells

The causal generative framework of scCausalVI enables robust generalization and cross-condition in silico prediction at single-cell resolution. Through manipulation of learned latent representations, the model predicts gene expression profiles under alternative conditions, allowing direct comparison within the same cellular context. This capability supports identifying treatment-induced differential expression and detecting the susceptible or resistant cells to interventions.

##### Identification of responsive cells in treated cohort

To distinguish treatment-induced changes from inherent variability, we develop a statistical framework comparing observed-to-counterfactual differences against a null distribution of uncertainty derived from the generative process. For each cell *x*_source_ from source condition, we generate both factual prediction 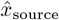, and cross-condition prediction of target condition, 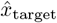, and project the concatenated dataset (observations and predictions) into a lower-dimensional PCA space. For simplicity, we adopt the same symbols to denote corresponding variables after PCA. Within this space, we formulate a permutation-based significance test: for the *i*th cell in source condition, let 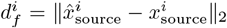 be factual difference, 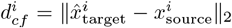 be the treatment-induced difference, with respective distributions *P*_*f*_ and *P*_*cf*_. We test the hypothesis:

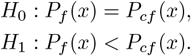

Cell *i* is classified as significantly responsive under *target* condition if 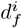 exceeds the (1 *− α*)-quantile of the empirical null distribution {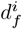: *i ∈* cells of source condition}, where *α* is the significance level. This non-parametric approach allows us to identify responsive cells while accounting for model uncertainty in the generative process.

### 4.5 Simulation procedure

We utilized scDesign3 [32] to simulate perturbed single-cell sequencing data by mimicking the distributions observed in real perturbed scRNA-seq data. scDesign3 utilizes a generalized additive model for location, scale, and shape (GAMLSS) to capture gene-specific marginal distributions as functions of cell states and design covariates, decomposing each gene’s marginal distribution into feature-specific intercepts, potential batch effects, condition effects, and cell-state-specific condition effects. In our study we generated realistic synthetic data based on an IFN-*β* stimulated scRNA-seq dataset [33]. For computational efficiency, we focused on B cells, CD4+ naive T cells, CD14+ monocytes, and activated T cells, and generated one group of control data as well as two groups of treated data. In the control group, we maintained cell state variations without introducing any condition effects. The first treatment group simulated a 1.5-fold condition effect exclusively on CD4+ naive T cells, while the second treatment group modeled a 2-fold condition effect solely on B cells. This design preserved intrinsic cell state variations while simulating differential cell-type specific responses to perturbations. To eliminate potential biases due to imbalances in conditions or cell types, we standardized the number of each cell type in each condition to 571 cells, which corresponds to the minimum number of cells across all cell types and conditions.

### 4.6 Datasets and pre-processing

We preprocessed all the scRNA-seq datasets using the standard workflow in the Scanpy package (version 1.9.6). Initially, raw count matrices were imported and cells with fewer than 50 detected genes or genes detected in fewer than 100 cells were filtered out to remove low-quality cells and genes. Then data normalization was performed by scaling each cell’s total counts to 10^6^ and applying a logarithmic transformation. Finally, 1,000 highly variable genes were identified using the ‘seurat v3’ method for downstream analysis by all the baseline models.

#### IFN-*β* data

We utilized the interferon-*β* stimulated single-cell RNA-seq dataset (GEO accession number GSE96583), which comprises 24,645 peripheral blood mononuclear cells (PBMCs). The dataset includes approximately equal numbers of cells from unstimulated and interferon-*β* stimulated conditions. In our analysis, cells from the unstimulated condition serve as the control group, while cells exposed to interferon-*β* are considered the treated group.

#### Respiratory epithelial COVID-19 data

We utilized a processed single-cell RNA-seq dataset with accompanying metadata available from the COVID-19 Cell Atlas (https://www.covid19cellatlas.org/index.patient.html). The raw count data can be downloaded from the Single Cell Portal: https://singlecell.broadinstitute.org/single_cell/study/SCP1289/. This dataset comprises 32,588 respiratory epithelial cells collected from healthy donors and COVID-19-infected patients. In our analysis, cells from healthy donors serve as the control group, while cells from infected patients are considered the treated group.

#### COVID-19 PBMC data from Blish et al

We analyzed a COVID-19 PBMC dataset from Blish et al. containing 44,721 cells from healthy donors and COVID–19–infected patients. Processed count matrices with de-identified metadata are available for download from the COVID-19 Cell Atlas hosted by the Wellcome Sanger Institute (https://www.covid19cellatlas.org/#wilk20). Additionally, the processed data are accessible for viewing and exploration on the publicly available cellxgene platform by the Chan Zuckerberg Initiative (https://cellxgene.cziscience.com/d/Single_cell_atlas_of_peripheral_immune_response_to_SARS_CoV_2_infection-25.cxg/). In our analysis, we designated cells from healthy donors as the control group and cells from infected patients as the treated group.

#### COVID-19 PBMC data from Meyer et al

To complement our batch-effect and negative-control analyses, we obtained a second COVID-19 PBMC dataset from Meyer et al., which can be downloaded from https://www.covid19cellatlas.org/index.patient.html. We selected only samples labeled “Adult” and restricted to the conditions “Healthy” or “COVID-19”, yielding a total of 67,383 cells. When performing batch-effect and negative-control validation, we focused on the cell types shared between the Blish and Meyer datasets for downstream integration, following our randomized experimental assumption. For batch-effect validation, healthy-donor cells served as the control group, whereas cells obtained from COVID-19-infected individuals were treated as the treatment group. In the integrative analysis, we used only healthy-donor cells from Blish and Meyer PBMC datasets, designating the Blish (healthy) cohort as the control and the Meyer (healthy) cohort as the treatment.

### 4.7 Quantitative metrics and baseline models

#### 4.7.1 Metrics

We used the average silhouette width (ASW)-based metrics and the Pearson correlation coefficient (PCC) for evaluation. The ASW-based metrics assessed the clustering or mixing quality of latent factors with respect to categorical labels. Specifically, ASW cond evaluated clustering based on condition labels, ASW celltype assessed clustering using cell type labels and ASW TE utilized affected labels in simulation data which indicated affected cells in treatment groups. A higher ASW score indicates better clustering of representations according to the given labels, while a score closer to zero suggests greater mixing regarding cell labels.

To measure batch mixing, we additionally employed the entropy score. For each cell, we computed the distribution of batch labels within its k-nearest-neighbor (k=30) neighborhood and calculated the corresponding entropy. A higher entropy score indicates more uniform (i.e., better) mixing of batches. For Harmony, we computed this entropy on the Harmony embedding; for scCausalVI, we calculated entropy on the background latent factors.

#### 4.7.2 Baseline models

We evaluated the performance of scCausalVI by benchmarking it against several state-of-the-art methods, including disentangled learning models (ContrastiveVI, scDisInFact, CPA, and biolord), a causal inference model (CINEMA-OT), and a generative model (scGen). To ensure a fair comparison, all methods were applied to the same datasets with identical preprocessing steps and parameter settings where applicable. For each method, we followed the recommended usage guidelines provided in their respective tutorials or application programming interfaces. Unless specified otherwise, default parameter settings were used. For deep learning-based models, we adjusted the dimensions of latent factors to ensure consistency across methods.

##### contrastiveVI

We used ContrastiveVI Python package (v0.2.0), which takes count data as input and models the data using a ZINB distribution. After preprocessing and normalization in Scanpy (v1.9.6), we initialized the model with two layers and set the dimensions of both latent spaces to 10 (*n salient latent=10, n background latent=10*, and *n layers=2*). We followed the usage instructions provided in the GitHub [tutorial](https://github.com/suinleelab/contrastiveVI). For the predicted data from ContrastiveVI, we applied normalization using *sc*.*pp*.*normalize total(pred, target sum=1e6* and log-transformation using *sc*.*pp*.*log1p(pred)* for downstream comparison.

##### scDisInFact

We utilized scDisInFact Python package (v0.1.0), which accepts count data as input and models it using a negative binomial (NB) distribution. Following the standard preprocessing steps, we initialized the model with both latent factor dimensions set to 10 (*Ks=[10, 10]*). The usage of scDisInFact followed the guidelines provided in the GitHub [tutorial](https://github.com/ZhangLabGT/scDisInFact/blob/main/demo.ipynb). Predictions from scDisInFact were normalized in the same manner as those from ContrastiveVI.

##### CINEMA-OT

We employed CINEMA-OT Python package (v0.0.5), which requires PCA-transformed data as input. After data preprocessing, we performed principal component analysis using sc.pp.pca(adata) on the log-normalized data and followed the instructions in the CINEMA-OT [tutorial](https://github.com/vandijklab/CINEMA-OT/blob/main/cinemaot_tutorial.ipynb).

##### scGen

We used scGen Python package (v2.1.1), which takes log-normalized data and cell type labels as input. Following data preprocessing, we adhered to the usage outlined in the scGen [tutorial](https://scgen.readthedocs.io/en/stable/tutorials/scgen_perturbation_prediction.html).

##### CPA

We implemented CPA Python package (v0.8.8), which accepts count data along with metadata such as cell type labels and dosage information. We assigned a dosage value of 1 to all treated cells. After data preprocessing, we initialized the model with default settings as per the CPA [tutorial](https://cpa-tools.readthedocs.io/en/latest/tutorials/), except we set *n hidden encoder=128, n layers encoder=2* and *n layers decoder=2*. Predictions from CPA were normalized following the same procedure as for ContrastiveVI.

##### biolord

We applied biolord Python package (v0.0.3), which requires log-normalized expression data and cell type labels as input. After data preprocessing, we followed the guidelines provided in the biolord [tutorial](https://biolord.readthedocs.io/en/latest/tutorials/biolord_pipeline.html), setting *decoder width=128, decoder depth=2, attribute nn width=128, attribute nn depth=2, n latent attribute categorical=10*.

##### Harmony

We implemented Harmony Python package (v0.0.10) by scanpy.external.pp.harmony integrate function after performing PCA. All parameters were kept at their default values as recommended in the Harmony documentation, allowing for batch correction across batches without altering the underlying biological variation. When comparing with the alignment of background latent factors by scCausalVI, the Harmony embedding was obtained by aligning cells across all four batch-condition combinations in the integrative analysis of Meyer and Blish datasets.

### 4.8 Code availability

scCausalVI is implemented as an open-source Python package and available at https://github.com/ShaokunAn/scCausalVI.

## 5 Acknowledgements

We acknowledge the support of the National Key Research and Development Program of China (NO. 2022YFA1004801 to L.W.). S.A. and M.H. were funded by NHGRI and a Data Insights grant from CZI.

J.C. was funded by the Helmsley foundation.

## 6 Author contributions

S.A. and L.W. conceived the study. S.A. implemented scCausalVI with the assistance of K.C. and J.X. S.A. performed computational analyses with the assistance of J.C. and M.H. S.A., L.W. and M.H. wrote the manuscript. L.W. and M.H. supervised the study.

## References

[1] Dixit, A. et al. Perturb-seq: dissecting molecular circuits with scalable single-cell rna profiling of pooled genetic screens. cell 167, 1853–1866 (2016).

[2] Gehring, J., Hwee Park, J., Chen, S., Thomson, M. & Pachter, L. Highly multiplexed single-cell rna-seq by dna oligonucleotide tagging of cellular proteins. Nature biotechnology 38, 35–38 (2020).

[3] Schiebinger, G. et al. Optimal-transport analysis of single-cell gene expression identifies developmental trajectories in reprogramming. Cell 176, 928–943 (2019).

[4] Peidli, S. et al. scperturb: harmonized single-cell perturbation data. Nature Methods 21, 531–540 (2024).

[5] Kramer, B. A., Sarabia del Castillo, J. & Pelkmans, L. Multimodal perception links cellular state to decision-making in single cells. Science 377, 642–648 (2022).

[6] Snijder, B. et al. Population context determines cell-to-cell variability in endocytosis and virus infection. Nature 461, 520–523 (2009).

[7] Lotfollahi, M., Wolf, F. A. & Theis, F. J. scgen predicts single-cell perturbation responses. Nature methods 16, 715–721 (2019).

[8] Lotfollahi, M. et al. Predicting cellular responses to complex perturbations in high-throughput screens. Molecular systems biology 19, e11517 (2023).

[9] Hetzel, L. et al. Predicting cellular responses to novel drug perturbations at a single-cell resolution. Advances in Neural Information Processing Systems 35, 26711–26722 (2022).

[10] Piran, Z., Cohen, N., Hoshen, Y. & Nitzan, M. Disentanglement of single-cell data with biolord. Nature Biotechnology 1–6 (2024).

[11] Lotfollahi, M., Naghipourfar, M., Theis, F. J. & Wolf, F. A. Conditional out-of-distribution generation for unpaired data using transfer vae. Bioinformatics 36, i610–i617 (2020).

[12] Weinberger, E., Lin, C. & Lee, S.-I. Isolating salient variations of interest in single-cell data with contrastivevi. Nature Methods 20, 1336–1345 (2023).

[13] Weinberger, E., Lopez, R., Hütter, J.-C. & Regev, A. Disentangling shared and group-specific variations in single-cell transcriptomics data with multigroupvi. In Machine Learning in Computational Biology, 16–32 (PMLR, 2022).

[14] Zhang, Z., Zhao, X., Bindra, M., Qiu, P. & Zhang, X. scdisinfact: disentangled learning for integration and prediction of multi-batch multi-condition single-cell rna-sequencing data. Nature Communications 15, 912 (2024).

[15] Träuble, F. et al. On disentangled representations learned from correlated data. In International conference on machine learning, 10401–10412 (PMLR, 2021).

[16] Dong, M. et al. Causal identification of single-cell experimental perturbation effects with cinema-ot. Nature Methods 20, 1769–1779 (2023).

[17] Bunne, C. et al. Learning single-cell perturbation responses using neural optimal transport. Nature methods 20, 1759–1768 (2023).

[18] Zhang, X.-H., Tee, L. Y., Wang, X.-G.Huang, Q.-S. & Yang, S.-H. Off-target effects in crispr/cas9-mediated genome engineering. Molecular Therapy-Nucleic Acids 4 (2015).

[19] Schölkopf, B. Causality for machine learning. In Probabilistic and causal inference: The works of Judea Pearl, 765–804 (2022).

[20] Yao, L. et al. A survey on causal inference. ACM Transactions on Knowledge Discovery from Data (TKDD) 15, 1–46 (2021).

[21] Imbens, G. W. & Rubin, D. B. Causal inference in statistics, social, and biomedical sciences (Cambridge university press, 2015).

[22] Pearl, J. Causality (Cambridge university press, 2009).

[23] Pawlowski, N., Coelho de Castro, D. & Glocker, B. Deep structural causal models for tractable counterfactual inference. Advances in neural information processing systems 33, 857–869 (2020).

[24] Johansson, F., Shalit, U. & Sontag, D. Learning representations for counterfactual inference. In International conference on machine learning, 3020–3029 (PMLR, 2016).

[25] Feuerriegel, S. et al. Causal machine learning for predicting treatment outcomes. Nature Medicine 30, 958–968 (2024).

[26] Zinati, Y., Takiddeen, A. & Emad, A. Groundgan: Grn-guided simulation of single-cell rna-seq data using causal generative adversarial networks. Nature Communications 15, 4055 (2024).

[27] Park, Y. P. & Kellis, M. Cocoa-diff: counterfactual inference for single-cell gene expression analysis. Genome Biology 22, 228 (2021).

[28] Lopez, R. et al. Learning causal representations of single cells via sparse mechanism shift modeling. In Conference on Causal Learning and Reasoning, 662–691 (PMLR, 2023).

[29] Bunne, C. et al. How to build the virtual cell with artificial intelligence: Priorities and opportunities. arXiv preprint 2409.11654 (2024).

[30] Hu, J., Shen, L. & Sun, G. Squeeze-and-excitation networks. In Proceedings of the IEEE conference on computer vision and pattern recognition, 7132–7141 (2018).

[31] Kiselev, V. Y., Andrews, T. S. & Hemberg, M. Challenges in unsupervised clustering of single-cell rna-seq data. Nature Reviews Genetics 20, 273–282 (2019).

[32] Song, D. et al. scdesign3 generates realistic in silico data for multimodal single-cell and spatial omics. Nature Biotechnology 42, 247–252 (2024).

[33] Kang, H. M. et al. Multiplexed droplet single-cell rna-sequencing using natural genetic variation. Nature biotechnology 36, 89–94 (2018).

[34] Hanahan, D. & Weinberg, R. A. Hallmarks of cancer: the next generation. cell 144, 646–674 (2011).

[35] Maan, H. et al. Characterizing the impacts of dataset imbalance on single-cell data integration. Nature Biotechnology 1–10 (2024).

[36] Zhang, Z. et al. Recovery of biological signals lost in single-cell batch integration with cellanova. Nature Biotechnology 1–17 (2024).

[37] Wilk, A. J. et al. A single-cell atlas of the peripheral immune response in patients with severe covid-19. Nature medicine 26, 1070–1076 (2020).

[38] Yoshida, M. et al. Local and systemic responses to sars-cov-2 infection in children and adults. Nature 602, 321–327 (2022).

[39] Korsunsky, I. et al. Fast, sensitive and accurate integration of single-cell data with harmony. Nature methods 16, 1289–1296 (2019).

[40] Goeva, A. et al. Hidden: a machine learning method for detection of disease-relevant populations in case-control single-cell transcriptomics data. Nature Communications 15, 9468 (2024).

[41] Ziegler, C. G. et al. Impaired local intrinsic immunity to sars-cov-2 infection in severe covid-19. Cell 184, 4713–4733 (2021).

[42] Zhou, Z. et al. Heightened innate immune responses in the respiratory tract of covid-19 patients. Cell host & microbe 27, 883–890 (2020).

[43] Schulte-Schrepping, J. et al. Severe covid-19 is marked by a dysregulated myeloid cell compartment. Cell 182, 1419–1440 (2020).

[44] Narasimhan, P. B., Marcovecchio, P., Hamers, A. A. & Hedrick, C. C. Nonclassical monocytes in health and disease. Annual review of immunology 37, 439–456 (2019).

[45] Hadjadj, J. et al. Impaired type i interferon activity and inflammatory responses in severe covid-19 patients. Science 369, 718–724 (2020).

[46] Chan, K. R. et al. Early peripheral blood mcemp1 and hla-dra expression predicts covid-19 prognosis. EBioMedicine 89 (2023).

[47] Oh, C.-k. et al. Targeted protein s-nitrosylation of ace2 inhibits sars-cov-2 infection. Nature chemical biology 19, 275–283 (2023).

[48] Wieczorek, M. et al. Major histocompatibility complex (mhc) class i and mhc class ii proteins: conformational plasticity in antigen presentation. Frontiers in immunology 8, 292 (2017).

[49] Sette, A. & Crotty, S. Adaptive immunity to sars-cov-2 and covid-19. Cell 184, 861–880 (2021).

[50] Zheng, M. et al. Functional exhaustion of antiviral lymphocytes in covid-19 patients. Cellular & molecular immunology 17, 533–535 (2020).

[51] Yap, J. K., Moriyama, M. & Iwasaki, A. Inflammasomes and pyroptosis as therapeutic targets for covid-19. The Journal of Immunology 205, 307–312 (2020).

[52] Komanduri, A., Wu, Y., Chen, F. & Wu, X. Learning causally disentangled representations via the principle of independent causal mechanisms. arXiv preprint 2306.01213 (2023).

[53] Khemakhem, I., Kingma, D., Monti, R. & Hyvarinen, A. Variational autoencoders and nonlinear ica: A unifying framework. In International conference on artificial intelligence and statistics, 2207–2217 (PMLR, 2020).

[54] Gasperini, M. et al. A genome-wide framework for mapping gene regulation via cellular genetic screens. Cell 176, 377–390 (2019).

[55] Rood, J. E., Hupalowska, A. & Regev, A. Toward a foundation model of causal cell and tissue biology with a perturbation cell and tissue atlas. Cell 187, 4520–4545 (2024).

[56] Kingma, D. P. & Welling, M. Auto-encoding variational bayes. arXiv preprint 1312.6114 (2013).

[57] Gretton, A., Borgwardt, K., Rasch, M., Schölkopf, B. & Smola, A. A kernel method for the two-sample-problem. Advances in neural information processing systems 19 (2006).

[58] Louizos, C., Swersky, K., Li, Y., Welling, M. & Zemel, R. The variational fair autoencoder. arXiv preprint 1511.00830 (2015).

