## supplementary figures for "scCausalVI disentangles single-cell perturbation responses with causality-aware generative model"

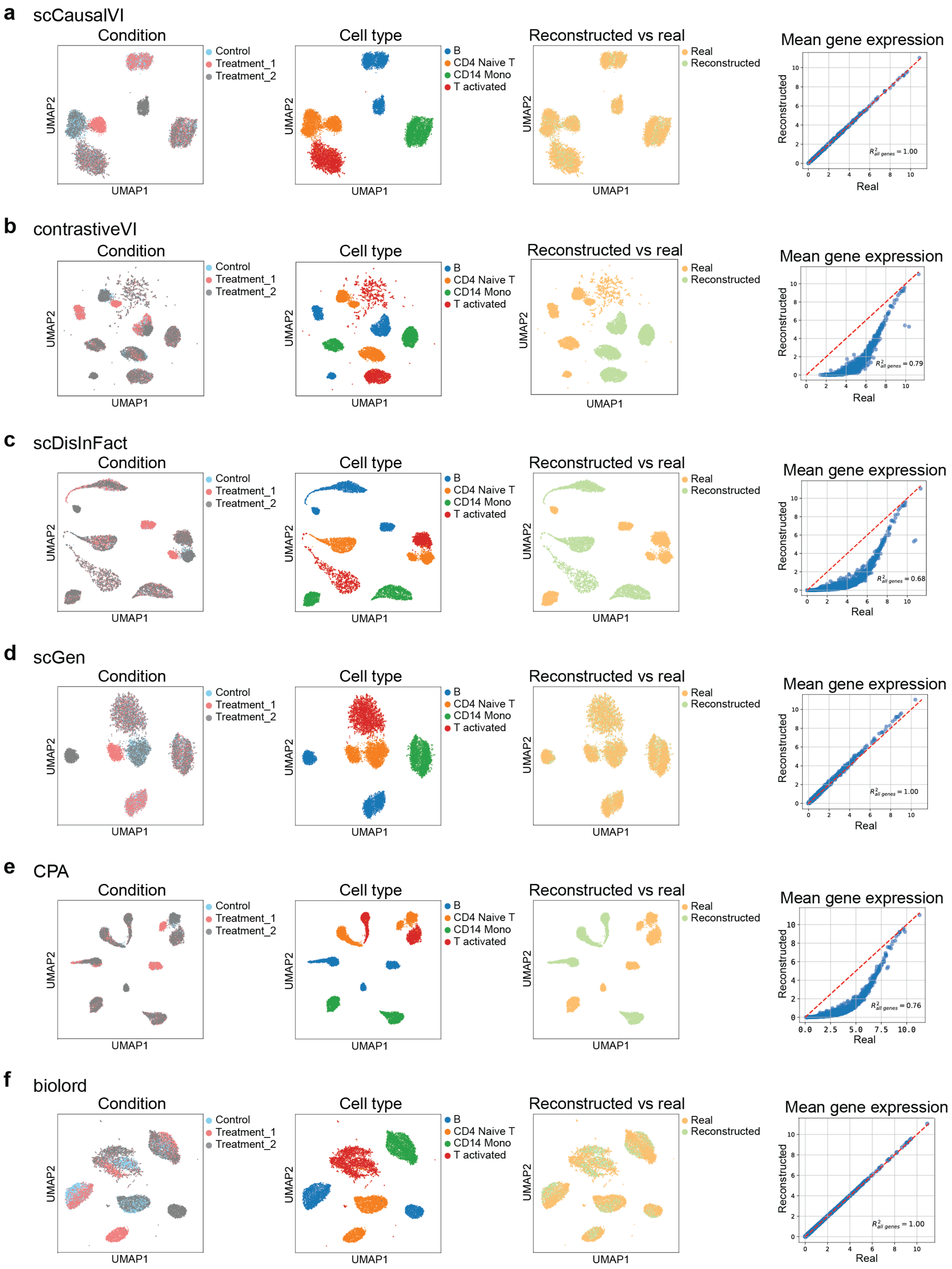


Supplementary Figure 1: **Prediction accuracy of generative models on simulated dataset. a-f**, UMAP visualizations of real and reconstructed data labeled by condition, cell type label, and data source, comparing the prediction accuracy for each generative model. Mean gene expression of all genes between reconstructed and real data ($R^2$ denotes squared Pearson correlation between ground truth and predicted values).

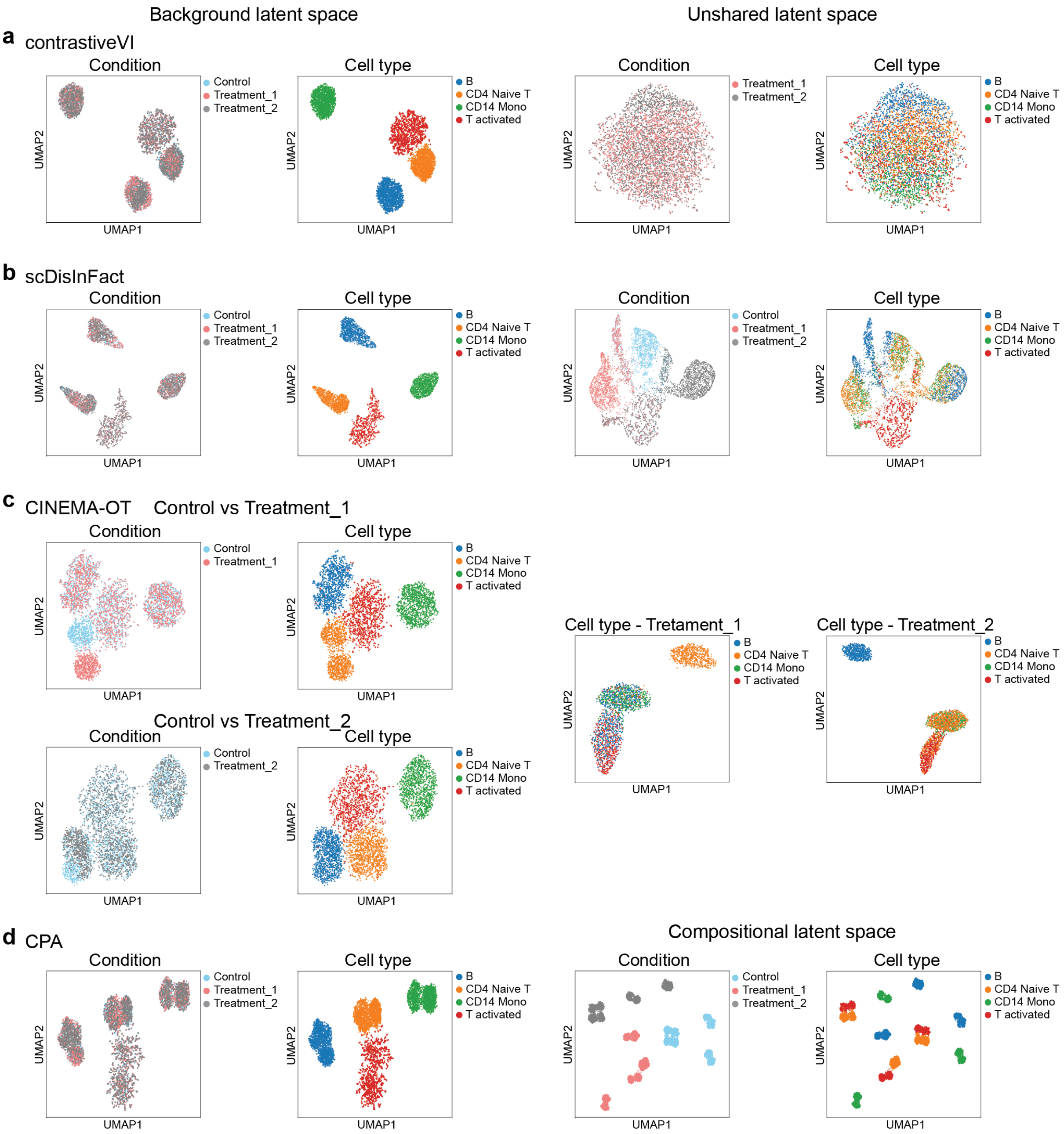


Supplementary Figure 2: **Disentanglement performance of baseline models on simulated dataset. a-c**, UMAP visualizations of latent factors, including background and condition-specific latent spaces, by disentangled methods—contrastiveVI **(a)**, scDisInFact **(b)**, and CINEMA-OT **(c)**. Due to its pair-wise optimal transport design, CINEMA-OT performs separate analyses for control vs Treatment_1 and control vs Treatment_2. **d**, UMAP visualization of basal latent factors of unperturbed cell states and compositional latent factors of perturbed variations by CPA.

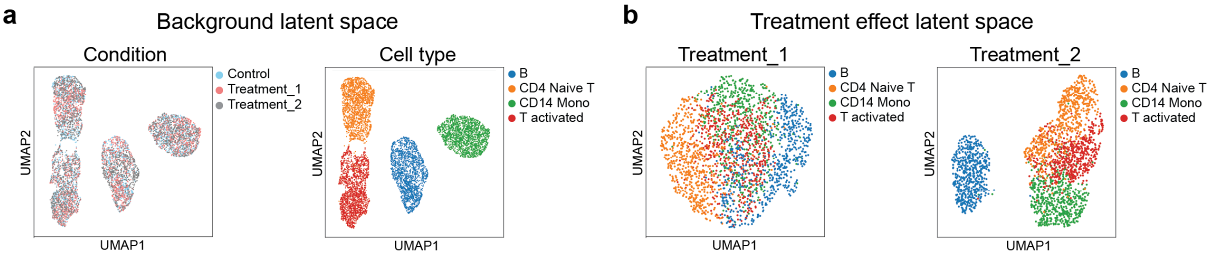


Supplementary Figure 3: **SENet ablation study on simulated data. a,** UMAP visualizations of background latent factors colored by condition and cell type label. **b,** UMAP visualizations of two sets of treatment effect latent factors colored cell type label.

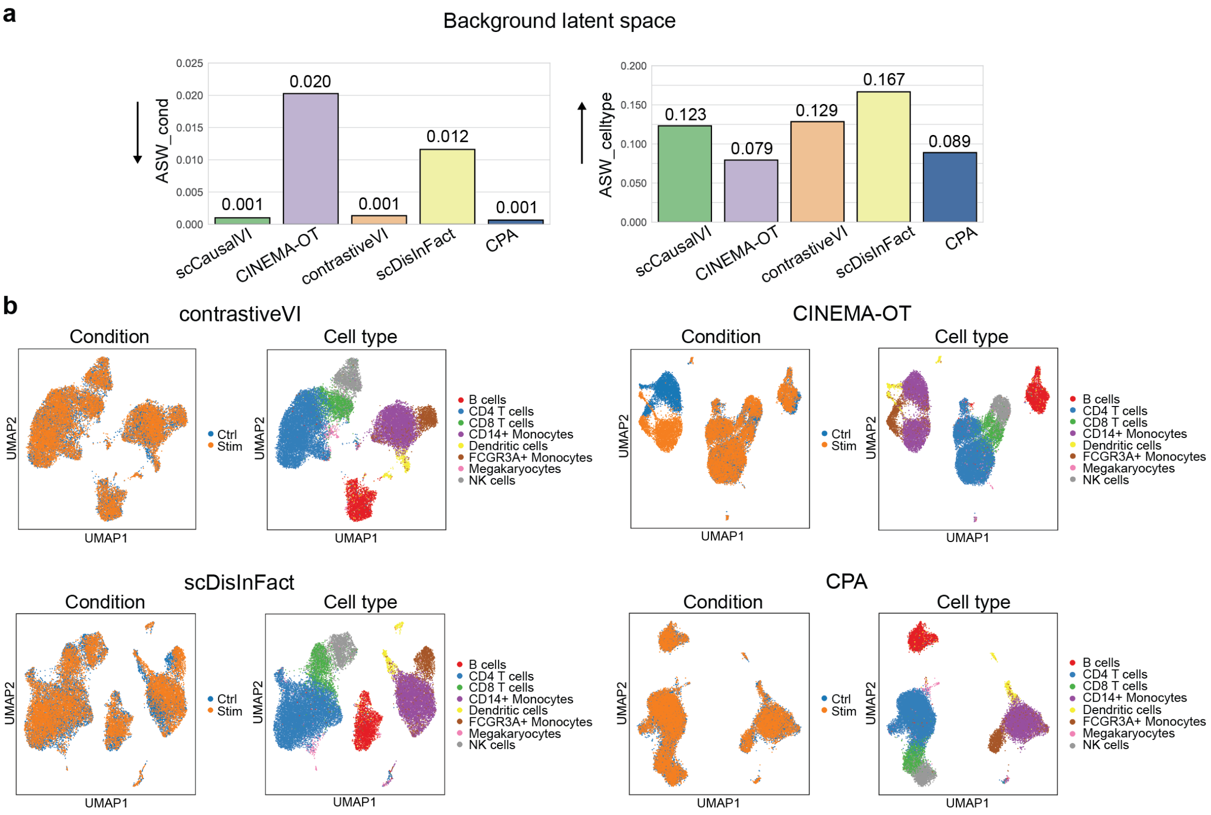


Supplementary Figure 4: **Cell type identity preservation in background latent space on IFN-$\beta$ dataset by baseline models a**, ASW-based metrics quantifying condition mixing and cell type clustering on background latent space of baseline models. **b**, UMAP visualizations of background latent factors by baselines, colored by conditions and cell type labels.

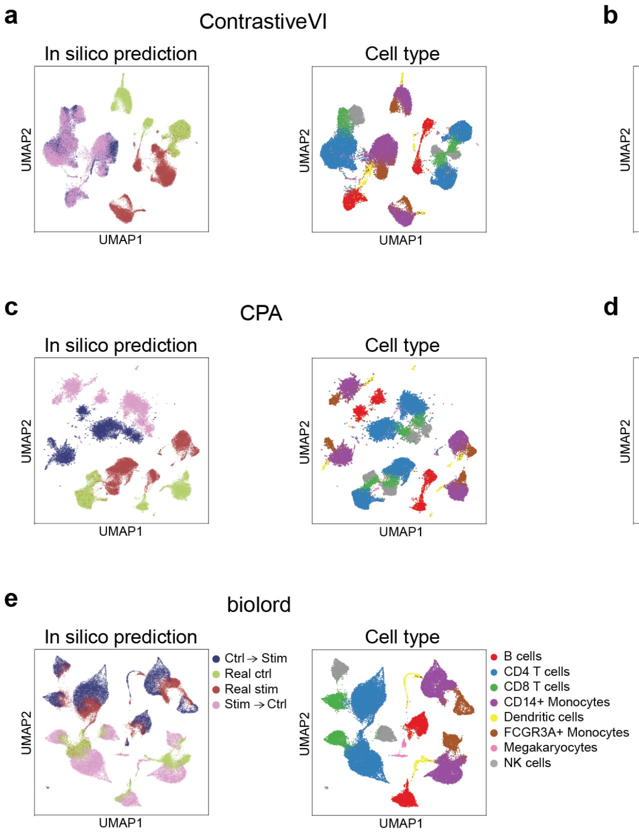


Supplementary Figure 5: **Cross-condition in silico prediction of baselines on IFN-$\beta$ dataset.** **a-e**, UMAP visualizations comparing real and cross-condition in silico prediction for baseline methods, colored by data source and cell type label.

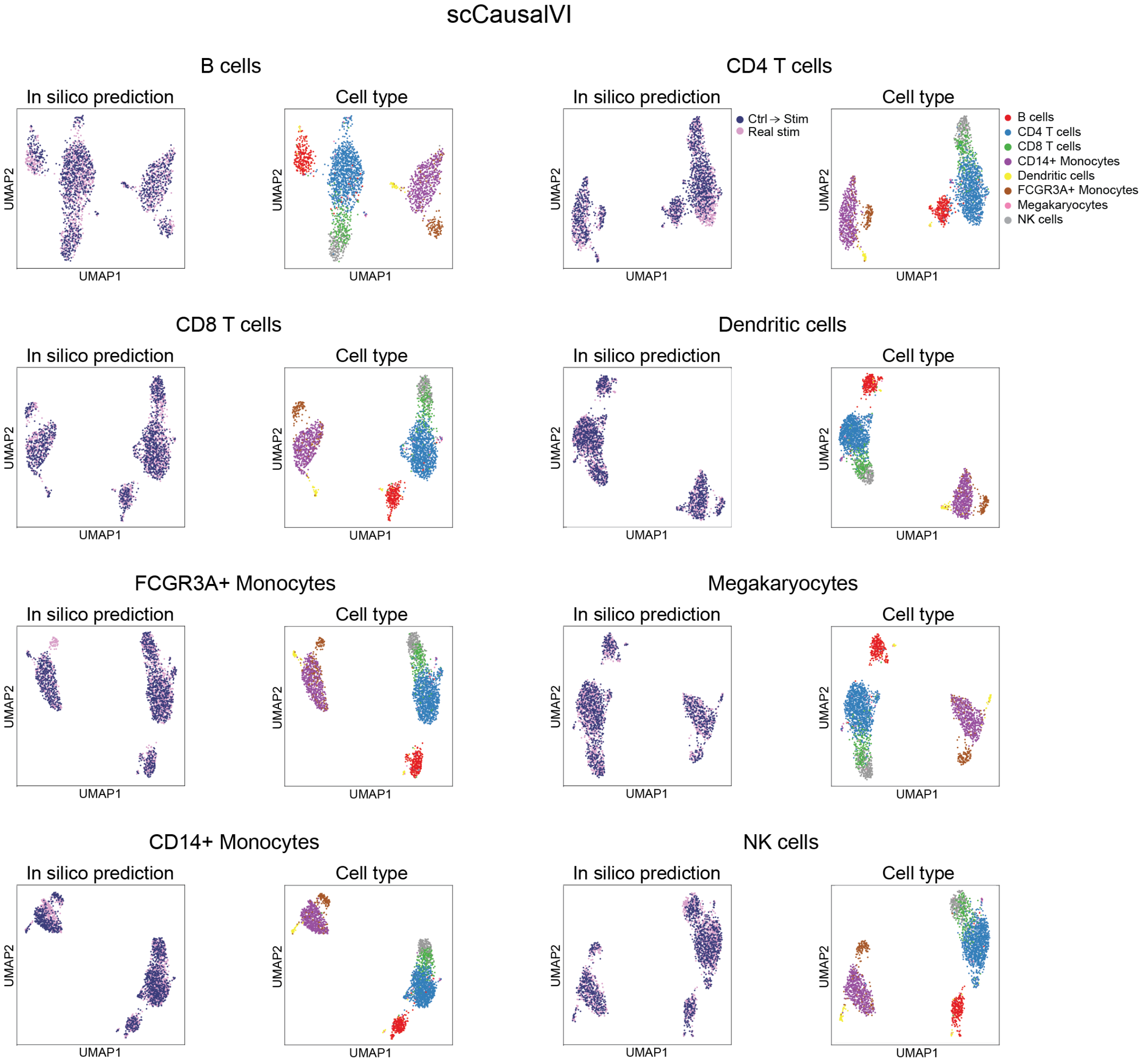


Supplementary Figure 6: **In silico prediction on OOD cells by scCausalVI with IFN-$\beta$ data**. UMAP visualizations comparing observed stimulated cells (Real stim) and cross-condition in silico predictions of control cells under stimulated conditions (Ctrl $\rightarrow$ Stim) by scCausalVI. For OOD generalization assessment, each cell type in train data was systematically withheld during model training and stimulated cohort of test data was used to assess prediction accuracy.

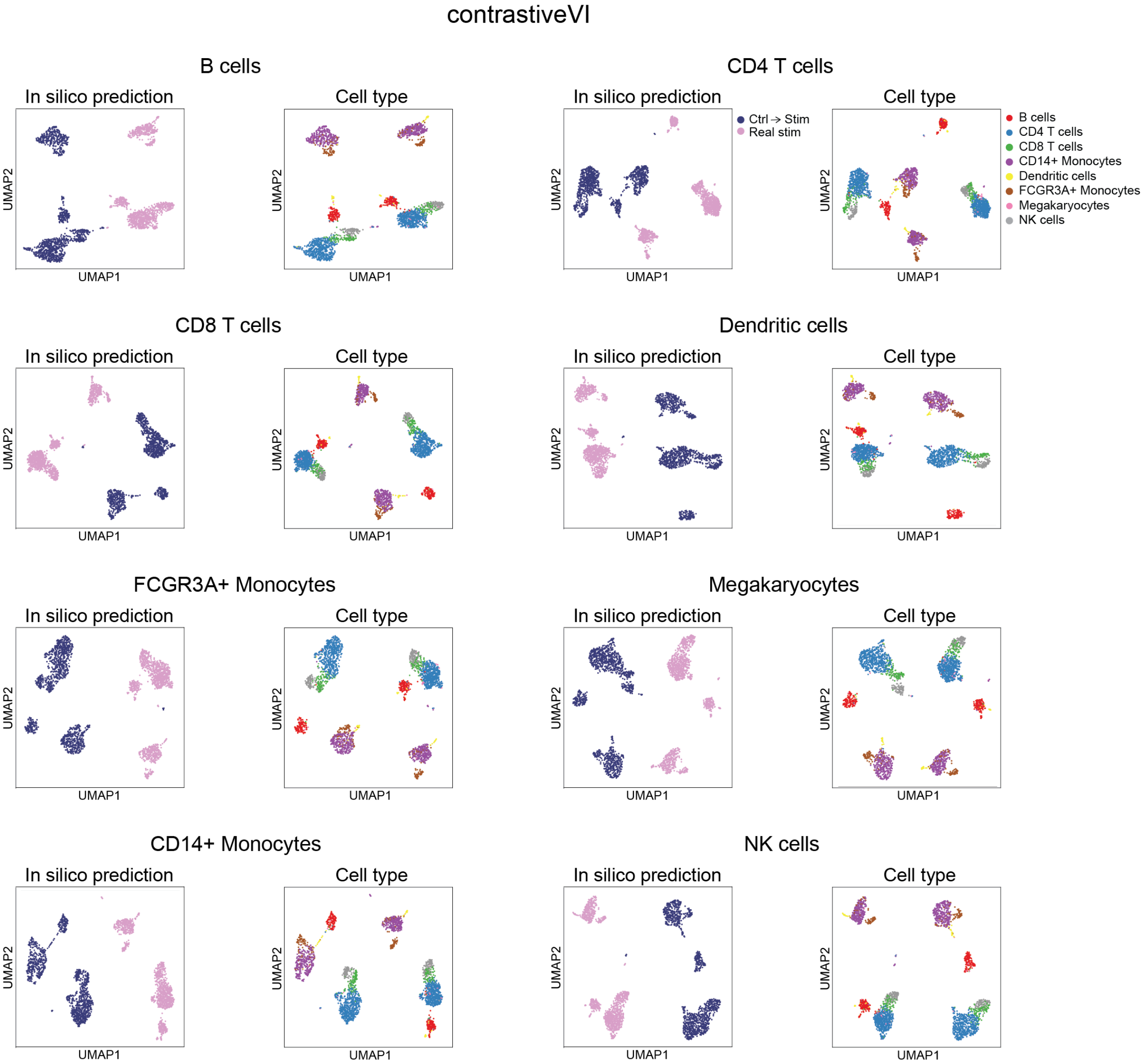


Supplementary Figure 7: **In silico prediction on OOD cells by contrastiveVI with IFN-$\beta$ data.** UMAP visualizations comparing observed stimulated cells (Real stim) and cross-condition in silico predictions of control cells under stimulated conditions (Ctrl $\rightarrow$ Stim) by contrastiveVI. For OOD generalization assessment, each cell type in train data was systematically withheld during model training and stimulated cohort of test data was used to assess prediction accuracy.

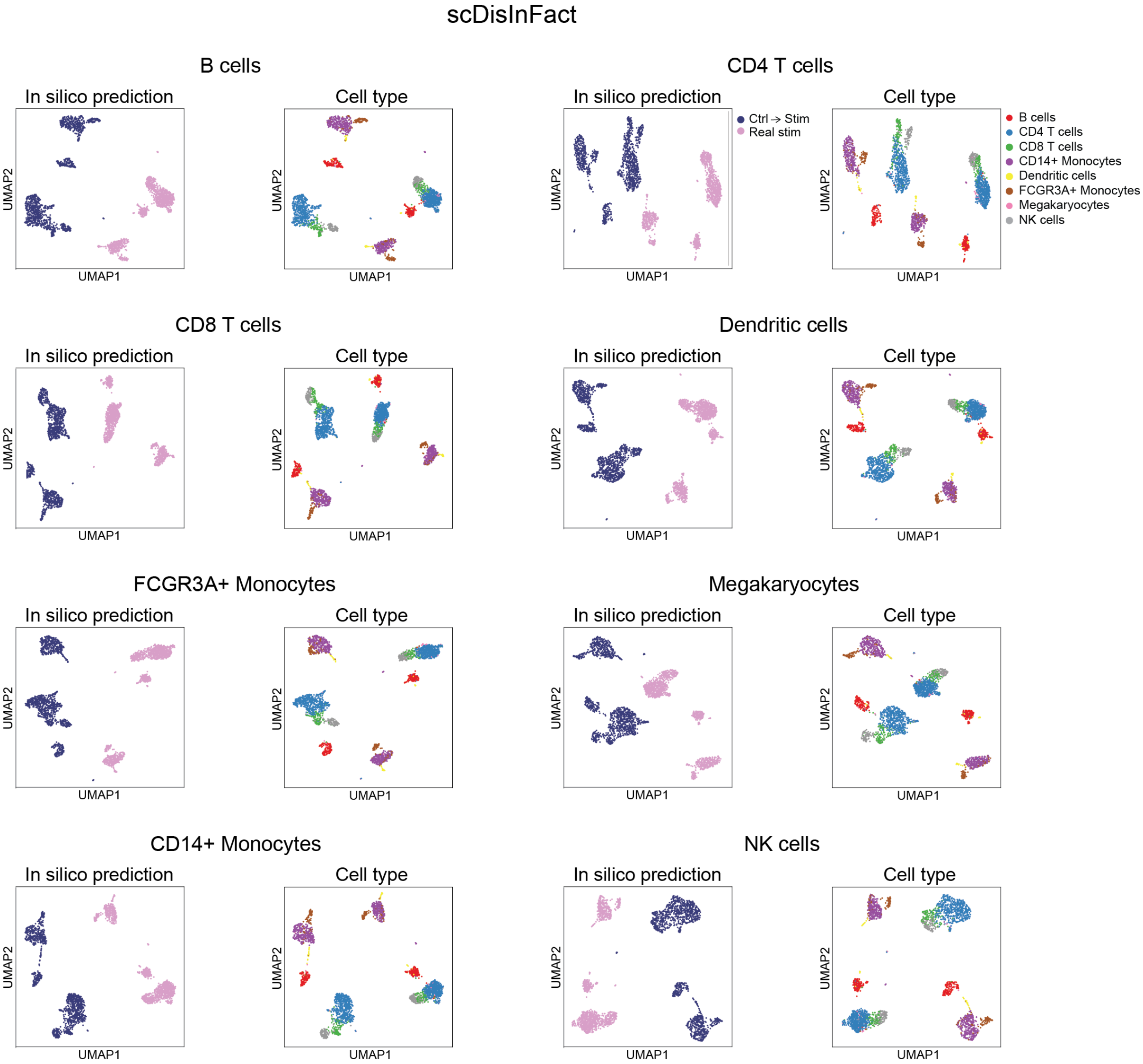


Supplementary Figure 8: **In silico prediction on OOD cells by scDisInFact with IFN-$\beta$ data**. UMAP visualizations comparing observed stimulated cells (Real stim) and cross-condition in silico predictions of control cells under stimulated conditions (Ctrl $\rightarrow$ Stim) by scDisInFact. For OOD generalization assessment, each cell type in train data was systematically withheld during model training and stimulated cohort of test data was used to assess prediction accuracy.

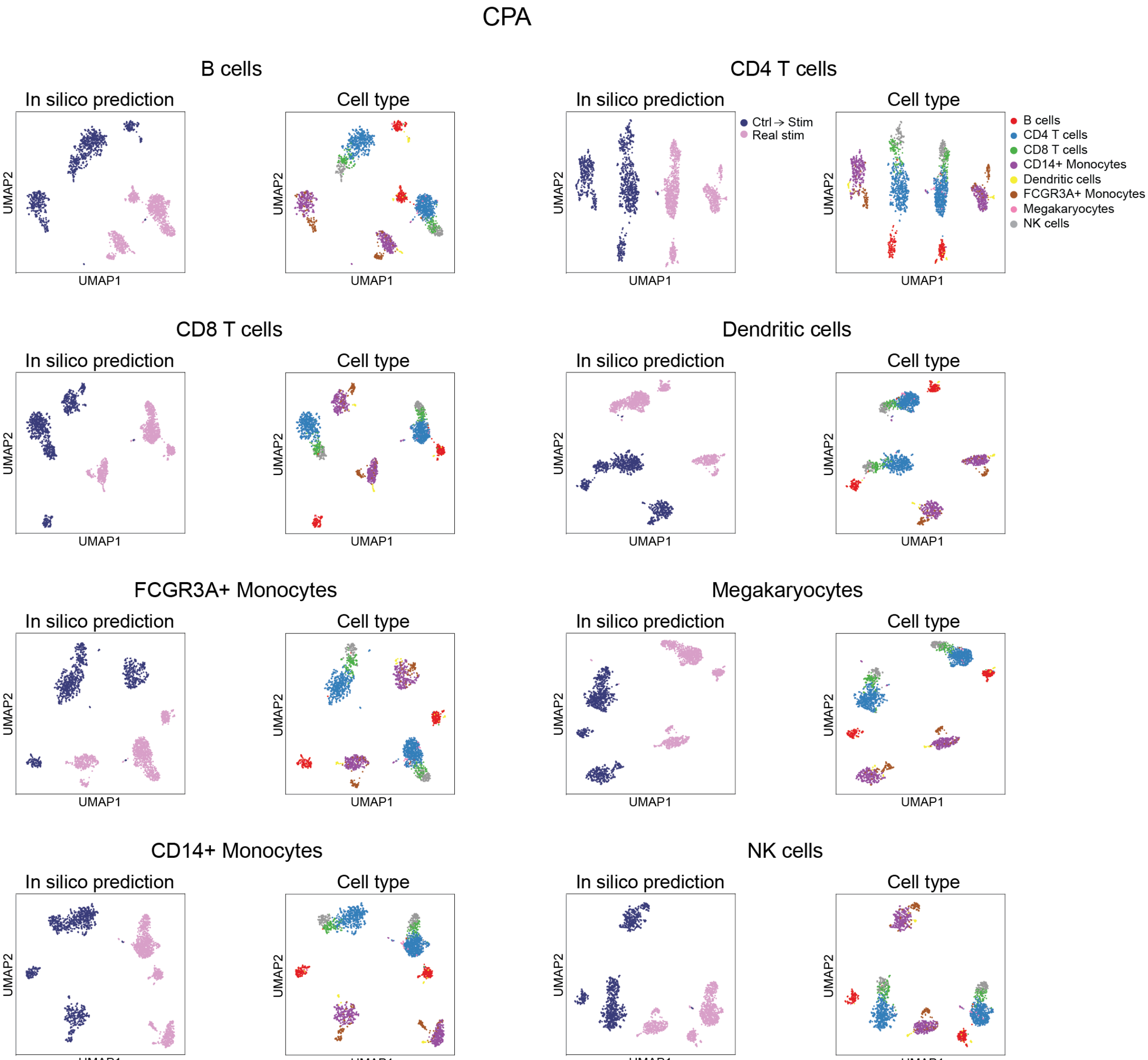


Supplementary Figure 9: **In silico prediction on OOD cells by CPA with IFN-$\beta$ data**. UMAP visualizations comparing observed stimulated cells (Real stim) and cross-condition in silico predictions of control cells under stimulated conditions (Ctrl $\rightarrow$ Stim) by CPA. For OOD generalization assessment, each cell type in train data was systematically withheld during model training and stimulated cohort of test data was used to assess prediction accuracy.

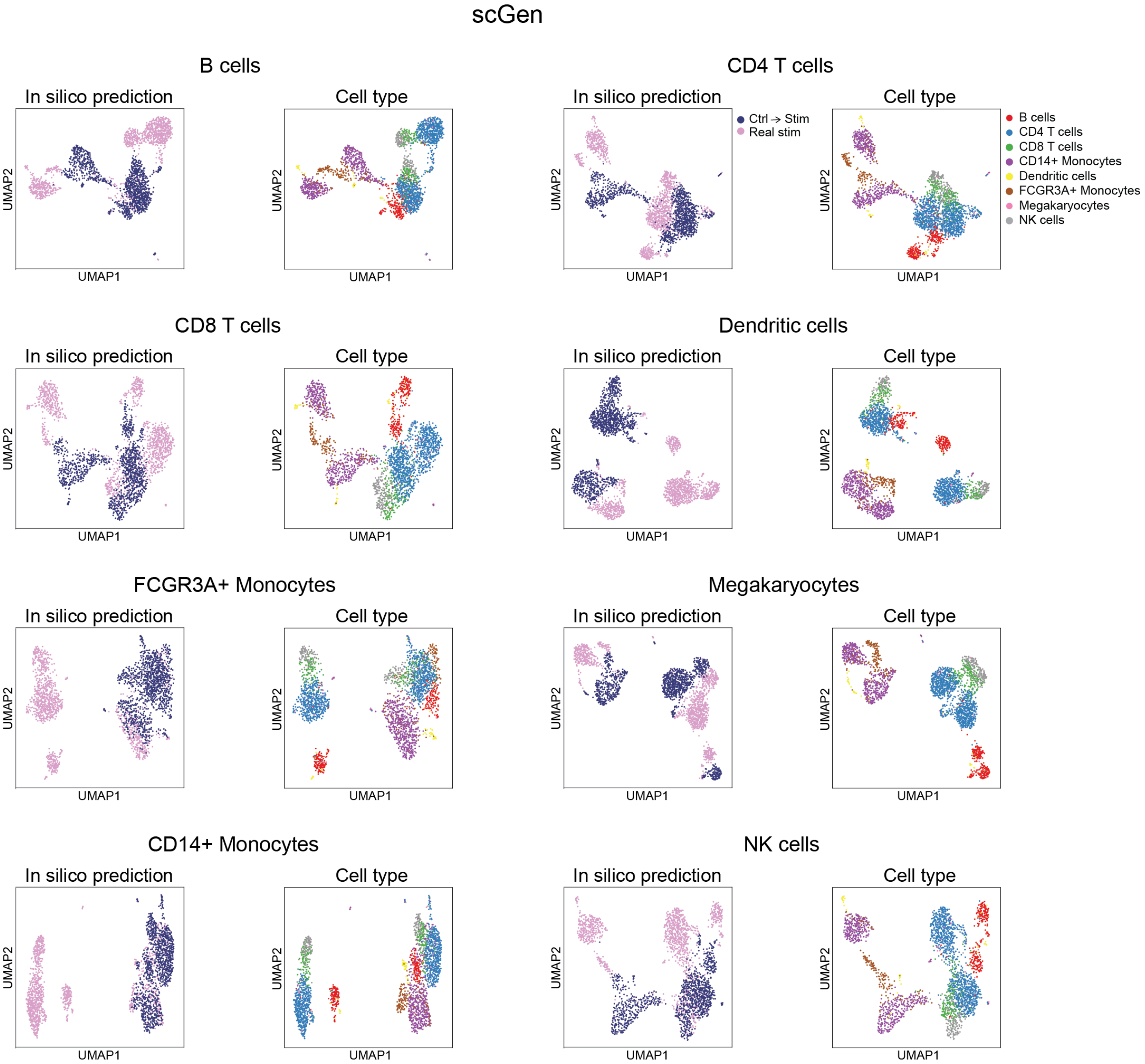


Supplementary Figure 10: **In silico prediction on OOD cells by scGen with IFN-$\beta$ data**. UMAP visualizations comparing observed stimulated cells (Real stim) and cross-condition in silico predictions of control cells under stimulated conditions (Ctrl $\rightarrow$ Stim) by scGen. For OOD generalization assessment, each cell type in train data was systematically withheld during model training and stimulated cohort of test data was used to assess prediction accuracy.

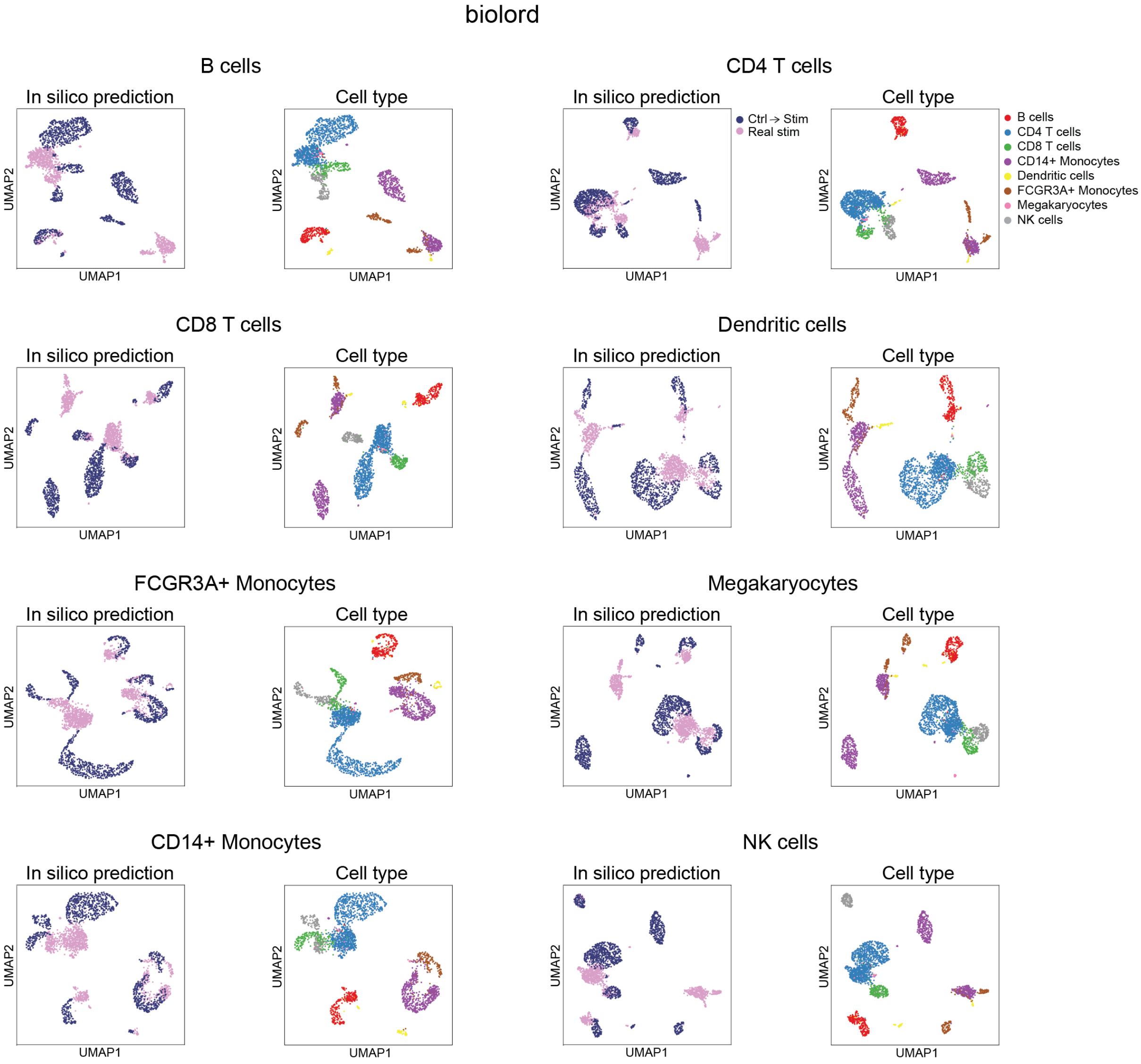


Supplementary Figure 11: **In silico prediction on OOD cells by biolord with IFN-$\beta$ data**. UMAP visualizations comparing observed stimulated cells (Real stim) and cross-condition in silico predictions of control cells under stimulated conditions (Ctrl $\rightarrow$ Stim) by biolord. For OOD generalization assessment, each cell type in train data was systematically withheld during model training and stimulated cohort of test data was used to assess prediction accuracy.

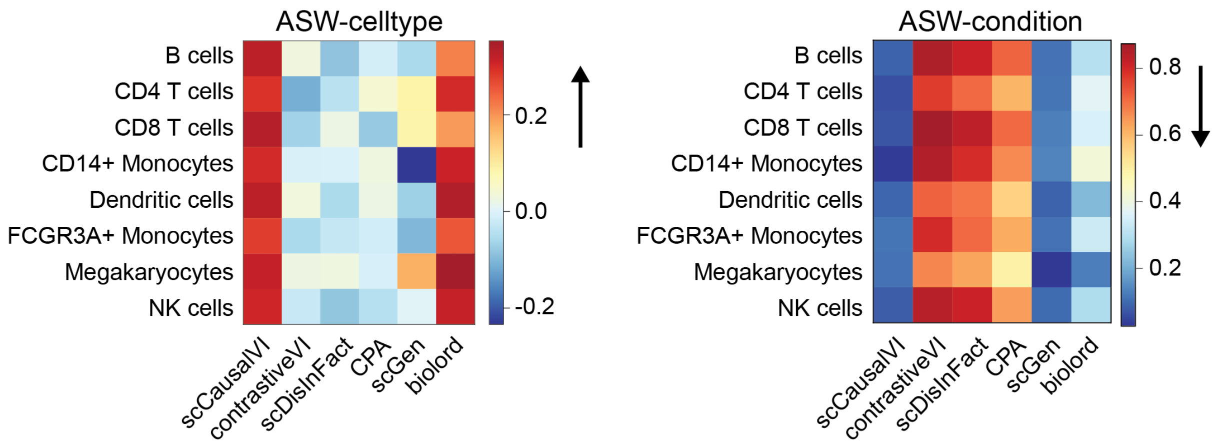


Supplementary Figure 12: **Cross-condition prediction performance on OOD IFN-$\beta$ data.** Heatmaps showing ASW-based metrics for cell type preservation (left) and condition mixing (right) between cross-condition predictions and observed stimulated OOD cells across different methods. For OOD evaluation, each cell type in train data was systematically excluded during model training and stimulated cohort in test data was used for evaluation. Metrics were computed using the top 50 principal components of concatenated predicted and observed data. The arrows indicate the direction of better performance.

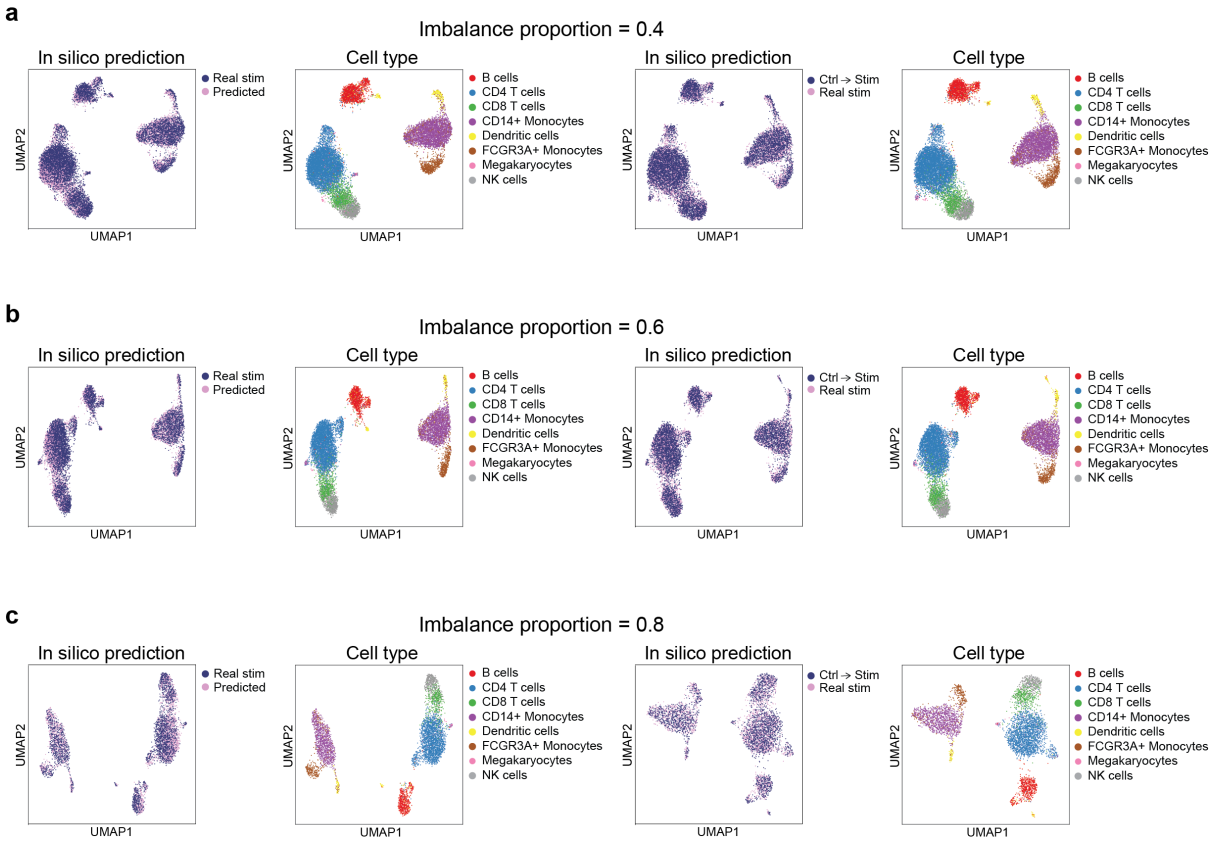


Supplementary Figure 13: **Robustness of scCausalVI under condition imbalance with IFN-$\beta$ data.** UMAP visualization comparing (from left to right): factual prediction accuracy between observed and predicted stimulated cells colored by data source, cell type identity, and cross-condition prediction accuracy colored by data source and cell type label. The stimulated cell population was systematically downsampled to (**a**) 40%, (**b**) 60%, and (**c**) 80% of its original size while maintaining the control group unchanged, to evaluate model performance under varying degrees of condition imbalance.

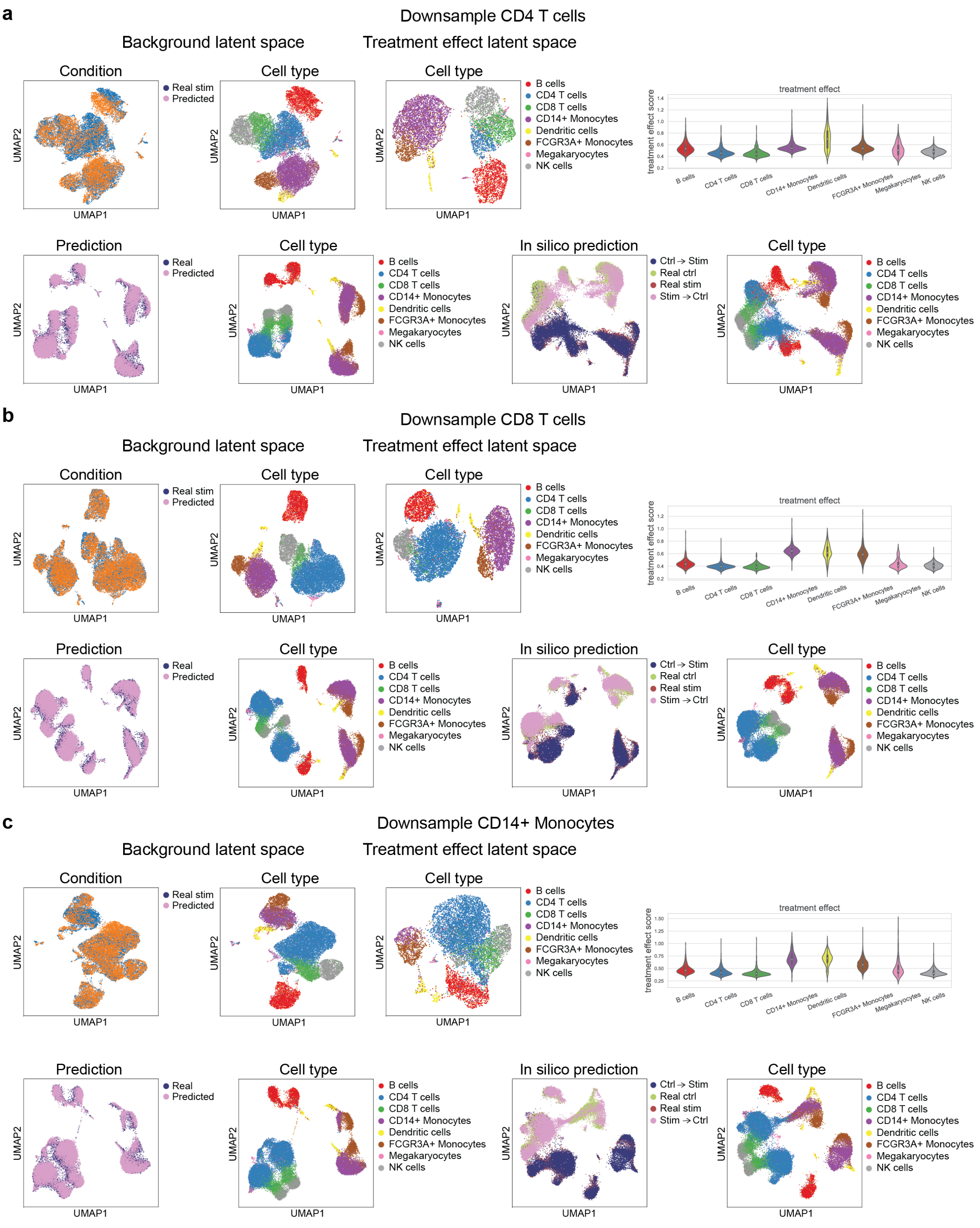


Supplementary Figure 14: **Robustness of scCausalVI under cell type imbalance with IFN-$\beta$ data**. For each panel in **a-c**, the upper row showed UMAP visualization of background latent space (colored by condition and cell type label), treatment effect latent space (colored by cell type label), and distribution of treatment effect scores across cell types. The lower row displayed factual prediction accuracy between observed and predicted data, and cross-condition prediction colored by data source and cell type label. During model training, cell populations in stimulated condition were selectively downsampled to 10% of original abundance for (**a**) CD4 T cells, (**b**) CD8 T cells, and (**c**) CD14+ Monocytes while maintaining other cell types unchanged.

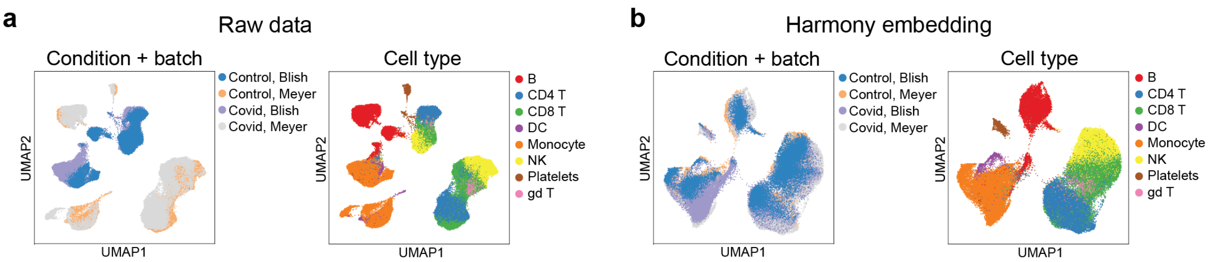


Supplementary Figure 15: **Visualization of batch effects in multi-batch COVID-19 datasets before and after batch correction by Harmony**. **a**, UMAP visualization of original data colored by condition and batch source (left) and cell type (right). **b**, UMAP visualization after Harmony batch correction colored by condition and batch source (left) and cell type labels (right).

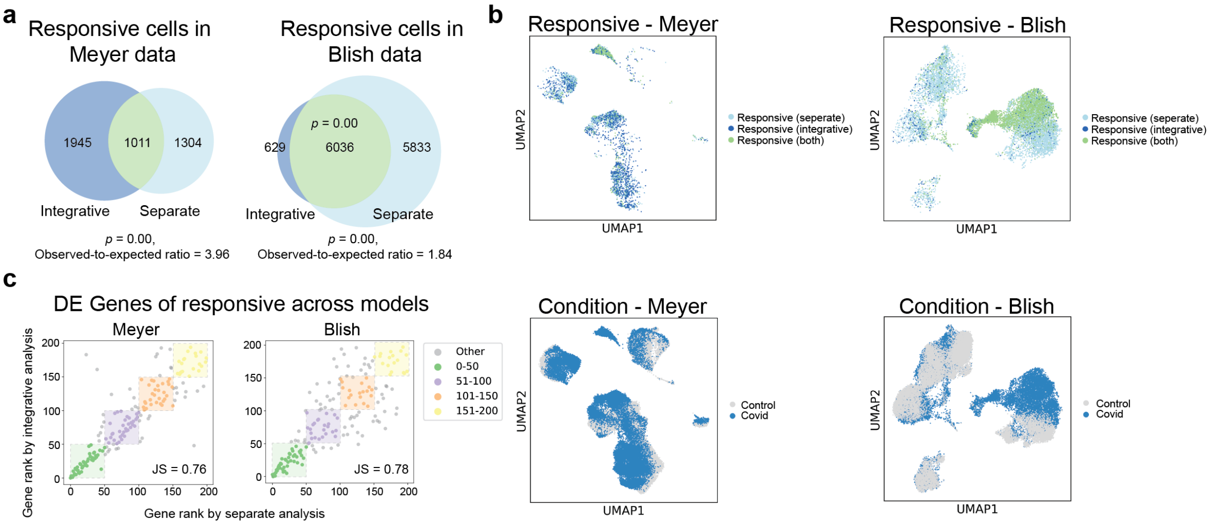


Supplementary Figure 16: **Comparison of integrative and separate analyses of Meyer and Blish datasets.** **a**, Venn diagrams showing the overlap of responsive cells identified through integrative and separate analyses in Meyer (left) and Blish (right) datasets. Hypergeometric tests showed significant overlapping cells ($p < 0.001$) with observed-to-expected ratios of 3.96 and 1.84, respectively. **b**, UMAP visualizations of responsive cells identified by integrative analysis (dark blue), separate analysis (light blue), or both methods (green) in Meyer (left) and Blish (right) datasets. **c**, Scatter plots comparing gene rankings of differentially expressed genes identified in responsive cells between integrative and separate analyses for Meyer (left) and Blish (right) datasets, with Jaccard similarity (JS) scores indicated. Colors denote genes grouped by their ranking positions (0-50, 51-100, 101-150, 151-200) in both analyses.

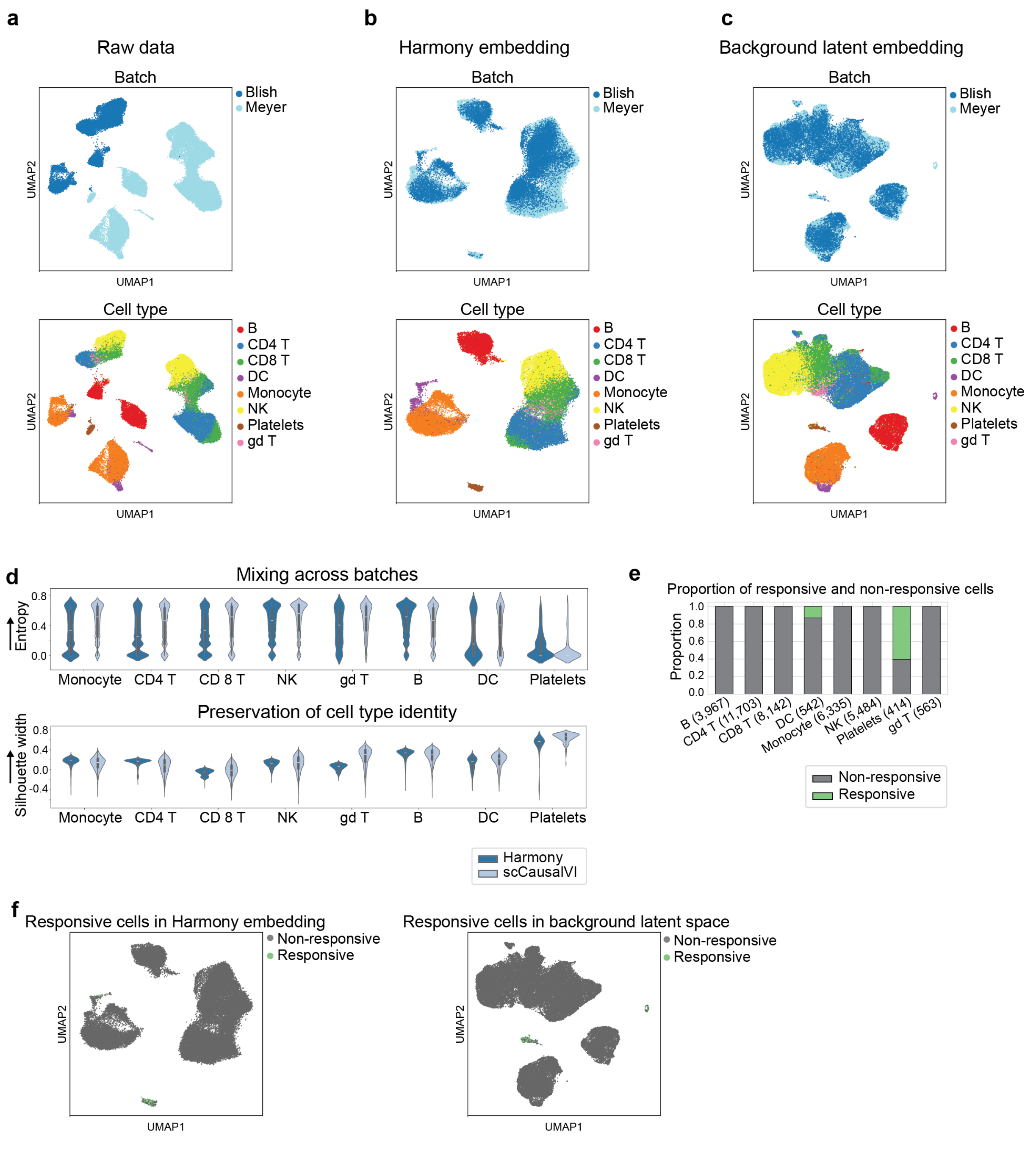


Supplementary Figure 17: **Negative control validation of scCausalVI using control samples from two independent batches of COVID-19 PBMC data.** **a-c**, UMAP visualization of raw data (**a**), Harmony-corrected embedding (**b**), and background latent factors by scCausalVI (**c**), from two independent COVID-19 PBMC datasets (Blish and Meyer), colored by batch source (top) and cell type labels (bottom). **d**, Violin plots comparing Harmony and scCausalVI performance across cell types for batch mixing (top) and cell type identity preservation (bottom). **e**, Bar plot showing the proportion of responsive and non-responsive cells across different cell types, with total cell counts shown in parentheses. **f**, UMAP visualizations of identified responsive cells in Harmony-corrected embedding (left) and in background latent space by scCausalVI (right).

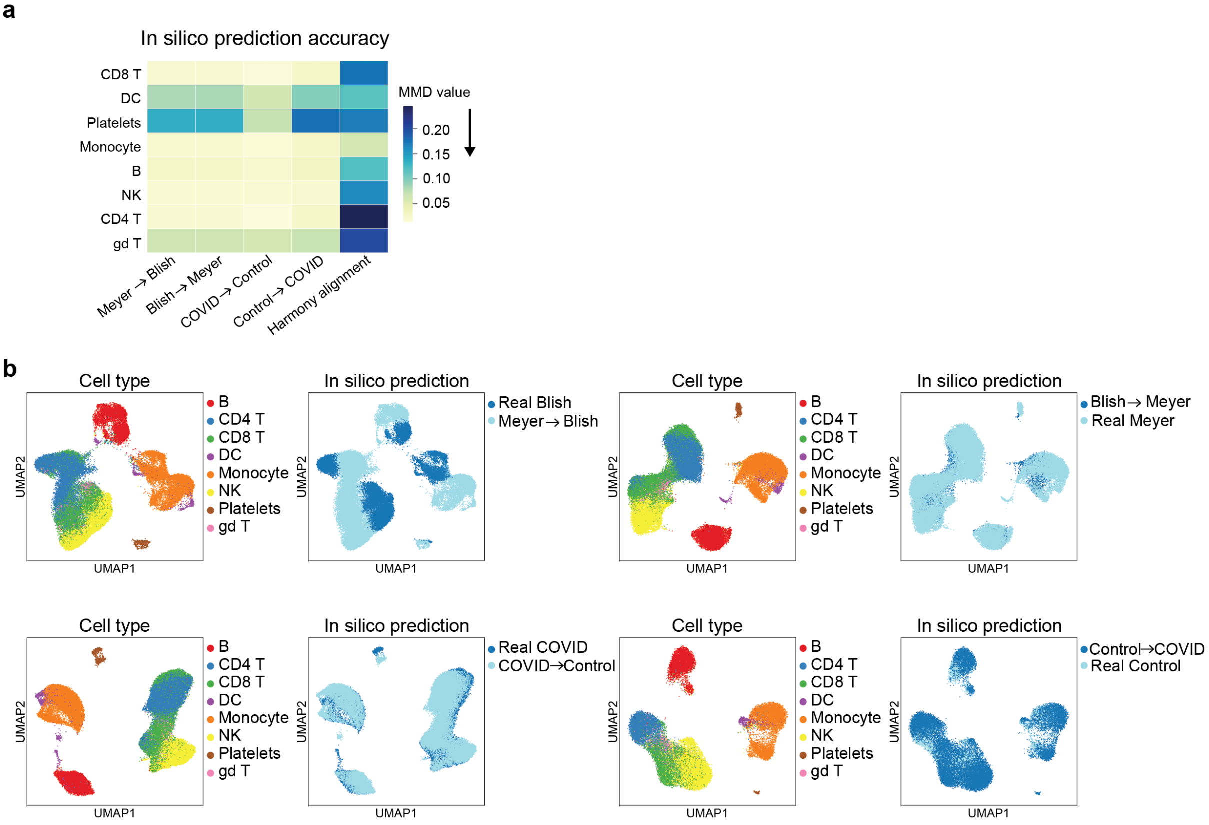


Supplementary Figure 18: **Negative control validation of scCausalVI by in silico prediction.** **a**, Heatmap of MMD values between in silico predictions and target populations calculated on the top 50 principal components of their joint PCA for both cross-batch and cross-condition predictions. A lower MMD indicates higher prediction accuracy. **b**, UMAP visualization of cross-batch and cross-condition in silico predictions and corresponding target populations, colored by cell type labels and conditions.

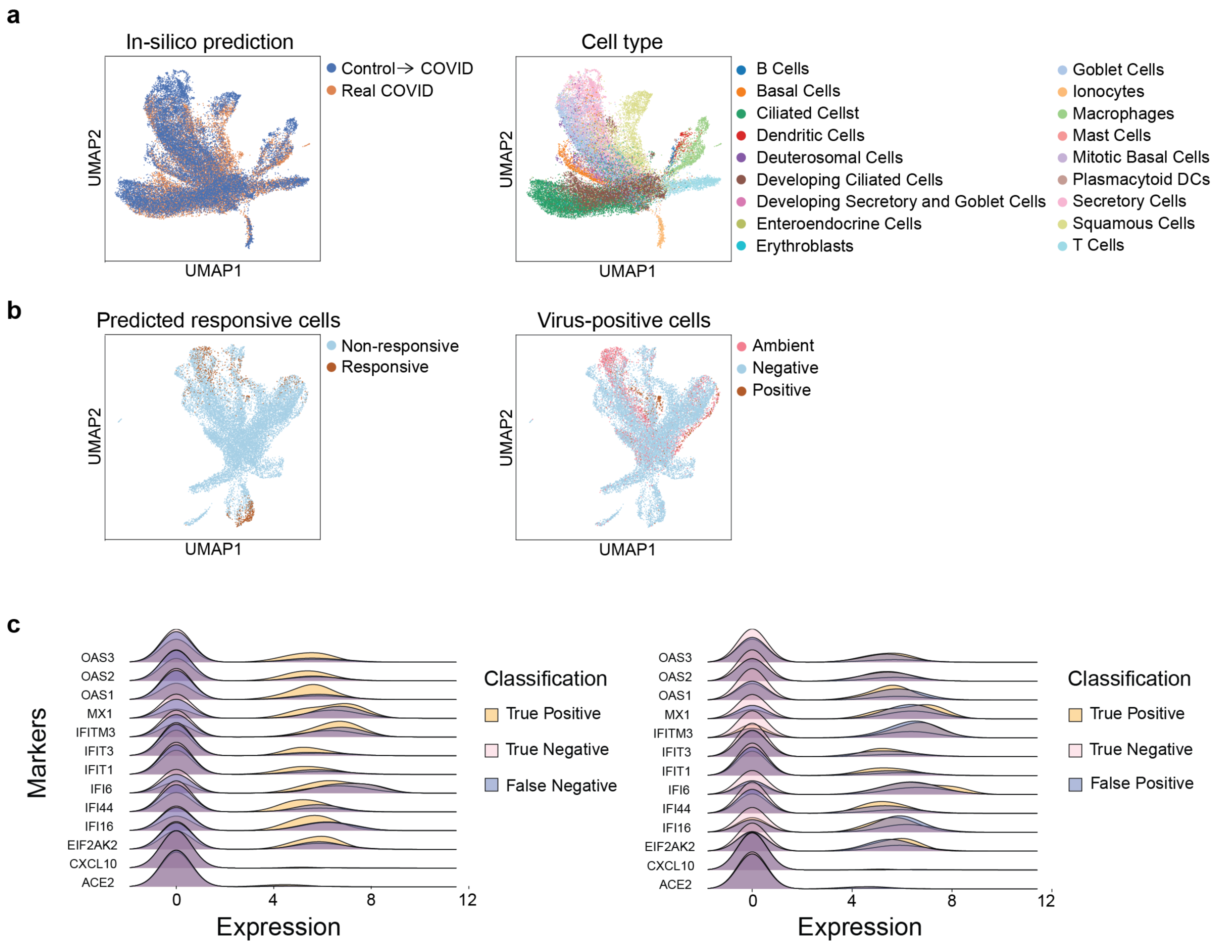


Supplementary Figure 19: **scCausalVI identified responsive and non-responsive respiratory epithelial cells in patient cohort**. **a**, UMAP visualization of real and in silico predictions, colored by data source and cell type labels. **b**, UMAP visualization pf cells from patients colored by predicted responsive labels (left) and viral transcripts detection labels (right). **c**, Distribution of normalized expression of interferon-stimulated genes and SARS-CoV-2 receptor, *ACE2*, classified by prediction results.

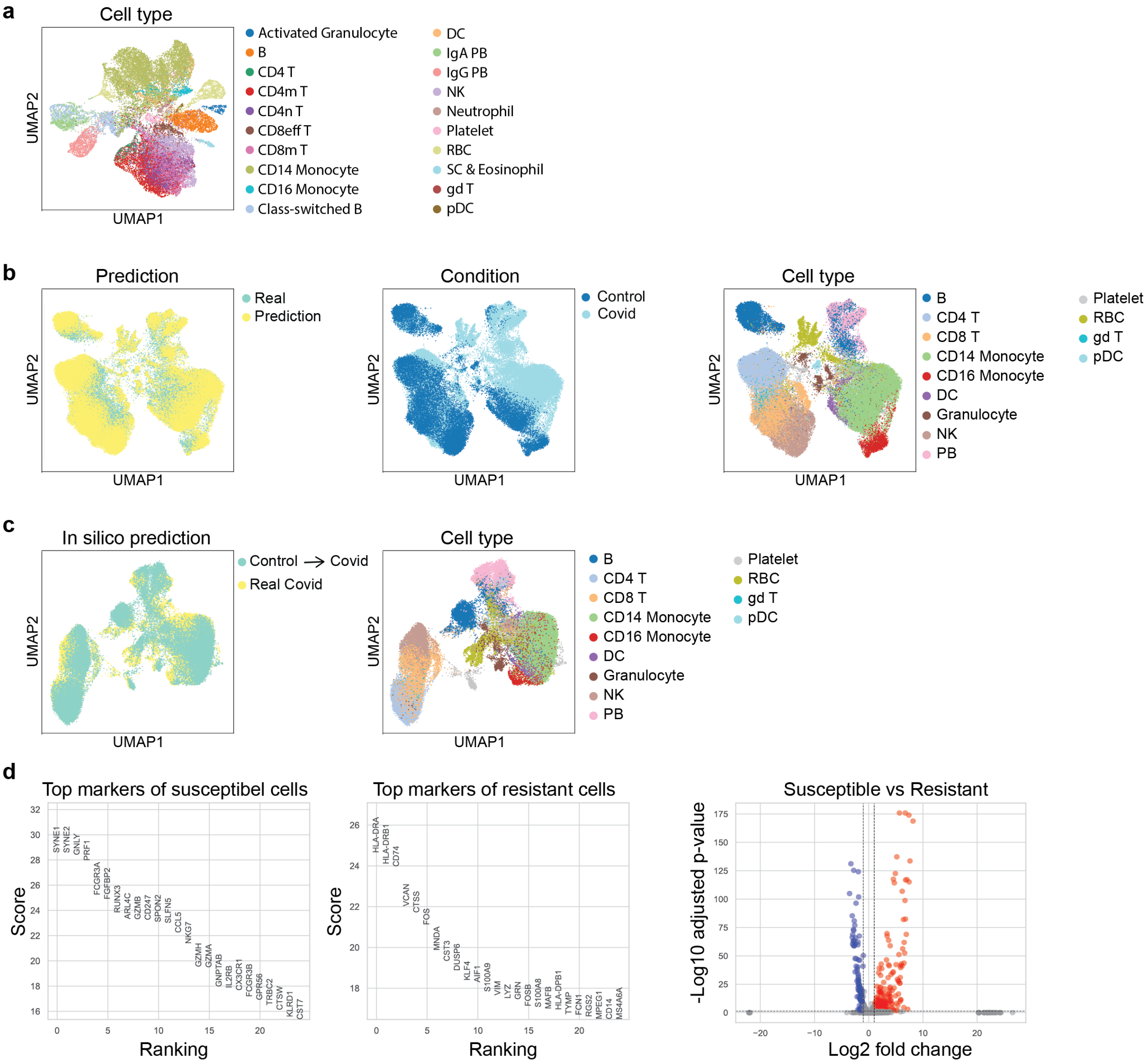


Supplementary Figure 20: **scCausalVI facilitated characteristics of susceptible and resistant cells by in silico prediction in PBMC COVID-19 data**. **a**, UMAP visualization of treatment effect latent factors colored by fine cell type labels. **b**, UMAP visualization of real and predicted data, colored by data sources, condition labels, and cell type labels. **c**, UMAP visualization of real and in silico prediction from patients, colored by data sources and cell type labels. **d**, Differential genes between susceptible and resistant cells.
